# Honeybee gut microbiota modulates host behaviors and neurological processes

**DOI:** 10.1101/2020.12.19.423587

**Authors:** Zijing Zhang, Xiaohuan Mu, Qina Cao, Yao Shi, Xiaosong Hu, Hao Zheng

**Affiliations:** College of Food Science and Nutritional Engineering, China Agricultural University, 100083 Beijing, China

**Keywords:** *Apis mellifera*, gut microbiota, social behavior, metabolism, gut-brain axis

## Abstract

Honeybee is a highly social insect with a reach behavioral repertoire and is a versatile model for neurobiological research. The honeybee gut microbiota is composed of a limited number of bacterial phylotypes that play an important role in host health. However, it remains unclear whether the microbiota can shape brain profiles and behaviors. Here, we revealed that the gut microbiota is requisite for the olfactory learning and memory ability of honeybees and alters the level of neurotransmitters in the brain. Transcriptomic and proteomic analysis showed distinctive gene expression and protein signatures for gnotobiotic bees associated with different gut bacteria. Specifically, genes related to olfactory functions and labor division are most upregulated. Moreover, differentially spliced genes in the brains of colonized bees largely overlapped with the datasets for human autism. The circulating metabolome profiles identified that different gut species regulated specific module of metabolites in the host hemolymph. Most altered metabolites are involved in the amino acid and glycerophospholipid metabolism pathways for the production of neuroactive compounds. Finally, antibiotic treatment disturbed the gut community and the nursing behavior of worker bees under field conditions. The brain transcripts and gut metabolism was also greatly interfered in treated bees. Collectively, we demonstrate that the gut microbiota regulates honeybee behaviors, brain gene transcription, and the circulating metabolism. Our findings highlight the contributions of honeybee gut microbes in the neurological processes with striking parallels to those found in other animals, thus providing a promising model to understand the host-microbe interactions via the gut-brain axis.

## Introduction

There is growing recognition that the gut microbiota plays a significant role in modulating host development and physiology, including metabolism and immune functions. Recent researches have focused on the effects of symbiotic microbes on the host’s central nervous system (CNS) and their involvement in the host behavioral processes. It reveals that gut microbiota can impact the host brain through a diverse set of pathways such as immune modulation and production of microbial metabolites implicated in the regulation of the gut-brain axis^1,2^. The symbiotic microorganisms inhabiting the gastrointestinal intestine are capable of producing various metabolites including neurotransmitters, amino acids, and short-chain fatty acids (SCFAs) that influence brain physiology^3-5^. While it is unclear if the neurotransmitters produced by certain gut bacteria (e.g., GABA, serotonin, and dopamine) can reach the brain considering their short half-lives and the block by the blood-brain barrier, the gut microbiota is capable of influencing brain physiology indirectly. Various SCFAs derived from microbial fermentation in the gut such as propionate and butyrate were suggested to regulate the rate-limiting enzymes involved in the biosynthesis of neurotransmitters in the brain^6^. Gut microbiota interferes CNS serotonergic neurotransmission by downregulating the level of tryptophan, the precursor of 5-hydroxytryptamine (5-HT), in the circulatory system, which further reduced the anxiety behavior in germ-free mice^5^. Furthermore, the expression level and alternative splicing of autism spectrum disorder (ASD)-related genes in the brain are proudly disturbed in mice harboring human ASD microbiome, which produces differential metabolome profiles^7^.

Although the significance of the functional connection between microbiota and neurophysiology has been widely appreciated, most current studies focused on the mammalian and non-social insect models. Further, it is challenging to unravel the distinctive contribution of individual gut members, which is partly due to the complex and erratic compositions of gut community and the difficulty to maintain and manipulate gnotobiotic animals^8^. Thus, models exhibiting high sociality and less complex gut community would be ideal to fully understand the relationship between the gut microbiota and host social behaviors.

Honeybee is a eusocial insect with distinct behavioral structures characterized by a complex range of interactive behaviors within the hive, and it has been extensively used as a model of perception, cognition, and social behaviors. A set of established methods are available to quantify the sophisticated behaviors of honeybee, such as associative appetitive learning and memory, sensory responsiveness, and hive behavioral observation^9^. It has been well documented that honeybees have a simple and host-specialized gut microbiota, with 8 ∼ 10 bacterial phylotypes comprising over 97% of the community^10-12^. Most bacterial phylotypes contain several divergent ‘sequence-discrete populations’ (SDPs) and a high extent of strain-level diversity^10,12^. All major bacterial phylotypes, including *Snodgrassella, Gilliamella, Bifidobacterium, Lactobacillus* Firm-4 and Firm-5, and *Bartonella* can be cultivated in the laboratory. Additionally, microbiota-free (MF) bees are experimentally tractable and can be colonized with defined communities of cultured strains^13,14^. Bee gut bacteria inhabit diverse niches and play specific roles in the bee gut, and they are beneficial to the host nutrition, immune homeostasis, and pathogen resistance^15^. These are probably accomplished via the microbial fermentation in the gut. The bee gut microbiota contributes to the degradation of diet polysaccharides, and untargeted metabolomics revealed that a plethora of organic acids accumulate in the presence of gut bacteria, which may have pivotal functional consequences in host physiology^13,16^.

Although the impact of gut community on the host’s health is relatively clear, few experiments have searched for the potential links between the honeybee gut microbiota and behavior. Pioneering explorations find that the level of biogenic amines (serotonin, dopamine, octopamine) implicated in bee behaviors is lower in newly emerged bees, which have an immature gut community^17^. In-lab-generated bees with a conventional (CV) gut microbiota behave differently in the gustatory responsiveness, and they possess altered endocrine signaling compared to the MF bees. Indeed, the gut microbiota affect the host metabolism that the hemolymph metabolomic profiles of CV and MF bees are separated^14^. The mono-association with the *Bifidobacterium asteroides* elevates the concentration of juvenile hormone III derivatives in the gut, which may regulate the host development^13^. All these findings strongly suggest that the honeybee gut microbiota may contribute to the host brain physiology and behavior phenotypes. Thus, it provides a particularly well-suited model to gain a detailed understanding of the gut microbiota-brain interactions.

Herein, we established gnotobiotic bees mono-colonized with different gut bacteria or with a conventional microbiota and identified that the presence of microbiota was sufficient to promote the host’s perception and cognition. Multi-omics analysis revealed that gut bacteria impact the neurotransmitter concentration, transcriptional program, protein level in the brain, as well as the circulating metabolic profiles. Finally, we confirmed that antibiotic exposure under field condition disturbs the hive behaviors of nurse bees, which is associated with altered brain transcripts and metabolite pools in the gut.

## Results

### Gut microbiota alters honeybee behaviors and brain neurotransmitter level

The ability to discriminate and memorize odors is critical for the social behaviors of honeybees, such as division of labor, organization of feeding, kin recognition, and mating^18,19^. We first examined whether the colonization of gut microbiota affects the olfactory learning and memory ability of bees under laboratory conditions. Each individual of CV, tetracycline treated (CV+tet), or MF bees generated in the lab (Methods, Supplementary Fig. 1a) was trained for 10 trials to associate the stimulus odor (nonanol) to a sucrose reward, and the memory test was performed 3h after the associative learning. Bees only responded to nonanol odor were considered to be a successful one (Fig. 1a, Supplementary Movie 1). Almost 50% of CV bees were able to memorize the nonanol odor and can distinguish the conditioned stimulus from the negative control odor (hexanol), and this percentage is similar to the previous test of hive bees performing the olfactory learning task^20^. In contrast, the proportion of successfully memorized individuals was significantly decreased in the antibiotic treatment group, and surprisingly, not an MF bee exhibited successful memory behavior (Fig. 1b). This suggests that the gut microbiota can apparently affect the learning and memory ability of bees. Proboscis extension response (PER) is a taste-related behavior that is fundamental for olfactory discrimination^21^. We then measured the PER of MF and CV bees, and bees mono-colonized with six different bacterial phylotypes (*Snodgrassella*, Sn; *Gilliamella*, Gi; *Bifidobacterium*, Bi; *Lactobacillus* Firm-4, F4; *Lactobacillus* Firm-5, F5; *Bartonella*, Ba) to estimate the olfactory sensation affected by the specific gut member. Compared to the MF group, CV bees are more sensitive to the low concentration of sucrose, which is consistent with the previous finding^14^. However, for the mono-colonized groups, only the colonization of F5 significantly elevated the sucrose sensitivity of bees (Fig. 1c), implying an integrative effect of the gut bacteria.

**Fig. 1.**
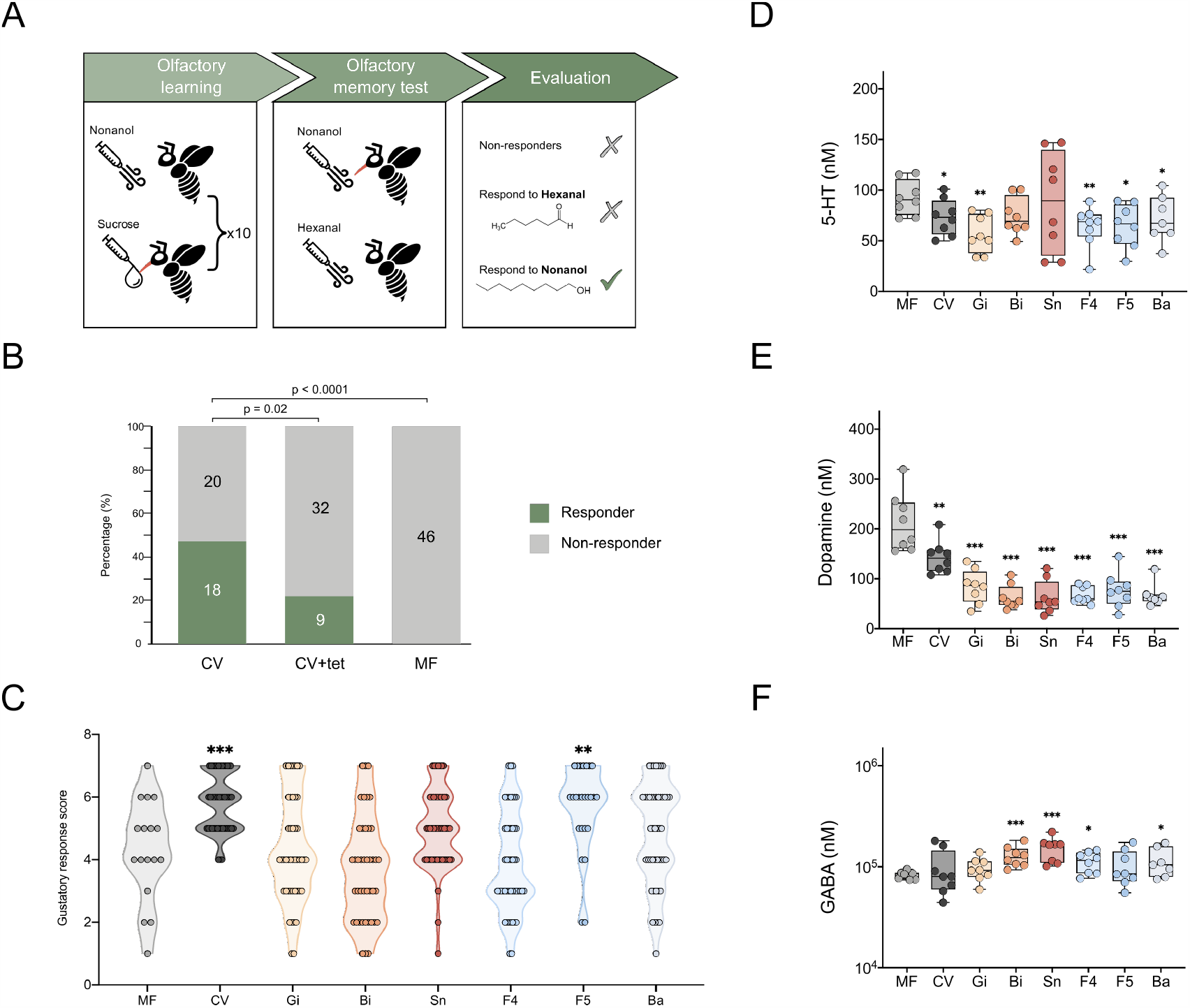
Gut microbiota alters honeybee behaviors and the concentration of neurotransmitters in the brain. (**a**) Olfactory learning and memory test design: 7-day-old conventionalized (CV), tetracycline-treated conventionalized (CV+tet), and microbiota-free (MF) bees were tested. Bees were trained to associate a conditional stimulus (nonanol odor) with a sucrose reward presented in ten successive trials. Bees responded only to nonanol odor in the memory test were considered successful. (**b**) Ratio of successfully memorized bees in the CV (n = 38), CV+tet (n = 41), and MF groups (n = 46). Group differences were tested by Chi-squared test. (**c**) Distribution of gustatory response score of MF (n = 17), CV bees (n = 46), and bees mono-colonized with different core gut bacteria: Gi, *Gilliamella apicola* (n = 45); Bi, *Bifidobacterium asteroides* (n = 42); Sn, *Snodgrassella alvi* (n = 50); F4, *Lactobacillus* Firm-4 (n = 48); F5, *Lactobacillus* Firm-5 (n = 25); Ba, *Bartonella apis* (n = 46). Each circle indicates a bee response to the provided concentration of sucrose. **p < 0.01, ***p < 0.001 (Mann–Whitney *u* test for the comparison with the MF group). (**d-f**) Concentrations of (**d**) 5-HT, (**e**) dopamine, and (**f**) GABA in the brains of MF (n = 8), CV (n = 8), and mono-colonized (n = 8, except n = 7 for Ba group) bees. Differences between bacteria-colonized and MF bees were tested by Mann-Whitney *u* test (*p < 0.1, **p < 0.01, ***p < 0.001). Error bars represent min and max (**d-f**).

Disordered olfactory behaviors are associated with alteration of neurotransmission in the bee brain^22,23^. Therefore, we investigated the changes in the brain’s neurochemistry of MF, CV, and mono-colonized bees. The concentration of five major neurotransmitters 5-HT, dopamine, GABA, tyramine, and octopamine that are important modulators of honeybee behaviors were determined in the brains. The concentration of 5-HT was significantly lower in CV bees and bees mono-colonized with Gi, F4, F5, and Ba than that in the MF bees (Fig. 1d). Likewise, dopamine that inhibits appetitive learning and decreases sucrose sensitivity in foragers^22,23^ was also decreased in bacteria-colonized bees (Fig. 1e). In contrast, the inhibitory transmitter GABA, which is required for fine odor discrimination^24^ and odor learning^25,26^, was significantly higher in the brains of Bi, Sn, and F4 bees (Fig. 1F). The biogenic amine, octopamine, and its precursor tyramine were not obviously altered by the conventional gut microbiota, while they are lowered in mono-colonized bee groups (Supplementary Fig. 1b, c). All these findings indicate that the colonization of either the normal gut microbiota or each single core gut member can affect the neurotransmitter levels in the brain, which might be associated with the altered olfactory sensitivity and learning-memory performance.

### Transcriptomic and alternative splicing profiles in the brain

The performance in PER and olfactory learning-memory behavior of honeybees is primarily associated with the gene expression profiles in the brain^27^. In total, our RNA sequencing analysis revealed that 713 genes were differentially expressed in bees colonized with gut members compared to MF bees (Supplementary Data 1), and different bee groups exhibited distinctive brain gene expression profiles (Supplementary Fig. 2a). Insect odorant-binding proteins (OBPs) play key roles in transport odorant molecules to olfactory receptors^28^, which is essential for the detection and distinguishment of specific odors^29^. Here, we found that the G protein-coupled olfactory receptor *Or115* and odorant binding protein *Obp14* were both upregulated in the CV group (Fig. 2a, Supplementary Fig. 2b), corroborating with the higher olfactory sensitivity of bees with a conventional microbiota (Fig. 1c). In addition, seven of the ten *mrjp* family genes of the major royal jelly protein (MRJP) encoding in *A. mellifera* genome were significantly upregulated in Bi and Sn groups, while bees colonized with *Gilliamella* exhibited decreased expression of the *mrjp* genes (Supplementary Fig. 2b). MRJPs have polyfunctional properties and participate in all major aspects of eusocial behavior in honeybees, such as caste determination and age polyethism^30^. Furthermore, genes encoding vitellogenin and the hexamerin HEX70a, which are both involved in the regulation of bee hormonal dynamics and the transition of foraging behavior^31,32^, were also upregulated in Bi and Sn groups (Supplementary Fig. 2b). The enrichment analysis of differentially expressed genes identified that KEGG pathways including linoleic, alpha-linolenic, arachidonic acids, and glycerophospholipid metabolism were upregulated in brains of different bacteria-colonized groups (Fig. 2b). The glycolysis/gluconeogenesis pathway that is critical for brain physiology via providing the fuel for brain functions^33^ was only upregulated in bees colonized with *Gilliamella*, while the protein processing, export, and ribosome pathways were upregulated in the Sn group. These results showed that the transcriptomic programs are differentially altered in the bacteria-colonized groups.

**Fig. 2.**
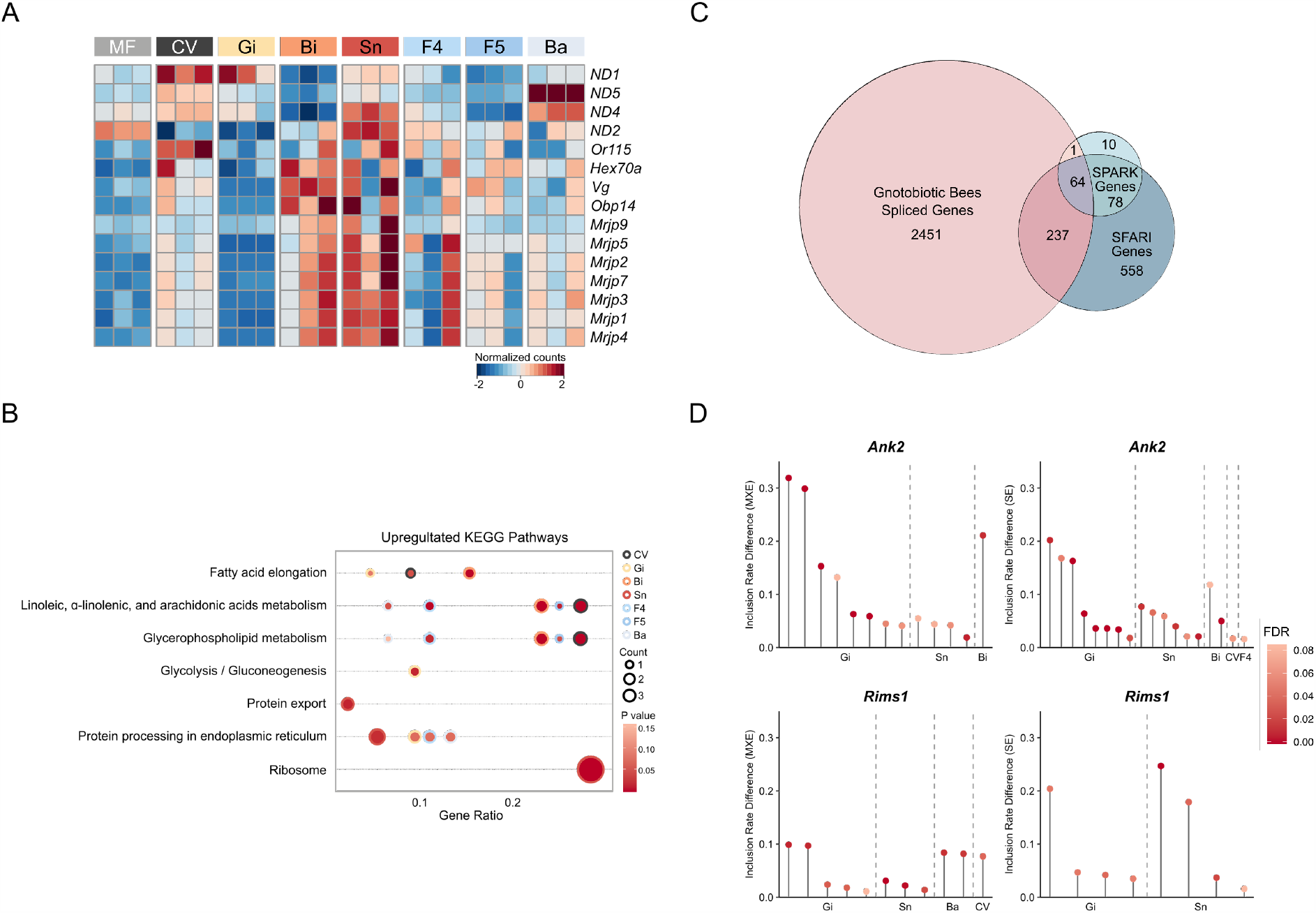
Gut microbiota impacts gene expression and alternative splicing of high confidence ASD genes in the honeybee brain. (**a**) Heatmap of differentially expressed genes in the brains of MF and bacteria-colonized bees. Each column represents one brain sample. Colors indicate the normalized gene counts. (**b**) KEGG pathways upregulated in the brains of CV or mono-colonized bees based on the differentially expressed genes. (**c**) Venn diagram of differentially spliced genes in the brains between MF and CV/mono-colonized bees (Gnotobiotic bees spliced genes; p < 0.05), and their overlap with the SPARK and SFARI Gene datasets. Differential splicing events were identified by rMATS. (**d**) Differentially splicing events (false discovered rate, FDR < 0.1) in *Ank2* and *Rims1* present in both SPARK and SFARI Gene datasets. Benjamini-Hochberg corrected p values (FDR) were calculated by rMATS. MXE, mutually exclusive exon; SE, skipped exon.

The gut microbiota does not only regulate gene expression but also affect alternative splicing (AS) of genes in the brain^7^, thus we investigate whether gut bacteria colonized bee brains show different AS events compared with the MF bees. rMATS analysis of the alternative splicing events of the brain genes detected a total of 22,064 events in 5,281 genes, and skipped exon (SE) is the most abundant among different types of AS. About 10–25% of events for each type of AS showed significantly different inclusion rates in bacteria-colonized bees (Supplementary Fig. 3a). The relative abundance of different types of AS events were similar across bee groups (Supplementary Fig. 3a). However, the UpSet plot shows that the vast majority of events do not intersect between sets, indicating that multiple AS events can occur in a single gene and the gut members cause different AS events (Supplementary Fig. 3b). Interestingly, it has been shown that the gene expression signatures of honeybees with disordered social behaviors are significantly enriched for human autism spectrum disordered (ASD)-related genes^34^. Likewise, the differentially expressed genes in bacteria-colonized bees also overlapped with those from human ASD patients (Supplementary Fig. 3c), implying the involvement of gut microbiota in host behaviors. Besides, dysregulation of alternative splicing in ASD-related genes is also associated with the psychiatric disorder^35^. Thus, we examined the overlap of genes showing significantly differential AS events between MF bees and bacteria-colonized groups with the ASD risk genes from the SPARK for Autism and the SFARI Gene datasets^36^ (Fig. 2c). Three hundred and two of the 2,753 differentially spliced genes in MF bees are associated with human autism, and 64 genes are present in both SPARK and SFARI Gene list (Fig. 2c). Interestingly, almost all identified homologs belong to the high-confidence SFARI gene list (Category 1) that have been clearly implicated in ASD (Supplementary Data 2). Specifically, we detected differential AS events of genes in MF bees in comparison with bacteria-colonized bees are related to the pathophysiology of ASD. For example, the inclusion rates of both mutually exclusive exon (MXE) and SE events in the *Ank2* gene that is important for neuronal migration^37,38^ are regulated in Gi, Sn, Bi, F4, and CV groups compared to MF bees (Fig. 2d). The synapse active-zone protein-coding gene *Rims1* with important roles in the maintenance of normal synaptic function^39^ also exhibited different inclusion rates of MXE and SE. Taken together, we identified that the gut microbes not only induce the differential gene expression profiles in the honeybee brain but also mediate AS resulting in specific gene isoforms. Genes essential for bee social behaviors and related to human ASD disease are apparently affected by different gut members, confirming the similarities in genes associated with social responsiveness of humans and honeybees^34^.

### Brain proteomics

Olfactory learning and memory behaviors of honeybees can be regulated by several proteins in the brain through proteomic analysis^40^. An in-depth proteome profile of the honeybee brain from MF and CV groups identified a total of 3,427 protein counts, 2,845 of which are both detected in MF and CV groups. Three hundred and twenty-two proteins were only found in MF bees, and 257 were exclusive for CV bees (Fig. 3a, Supplementary Data 3). Hierarchical cluster analysis of differentially expressed proteins shared by both groups demonstrated a clear separation between MF and CV bees (Fig. 3b). Notably, the muscarinic acetylcholine receptor (mAChR) involved in the cholinergic neurotransmitter system is upregulated in CV bees. mAChR is an acetylcholine binding receptor processing olfactory signals and plays an important role in the retrieval process of associative and non-associative learning and the formation of memory^41^, corroborating our findings of increased memory ability for CV bees (Fig. 1b). We also identified that a splicing factor U2af28 was upregulated in CV brains, supporting the differential patterns of AS in the brain. Interestingly, GO enrichment analysis of unique protein in the CV brain identified that GO terms are related to synaptic neurotransmission and cation/ion transmembrane transportation (Fig. 3c), which are essential for the fundamental functions in the honeybee central nervous system^42^.

**Fig. 3.**
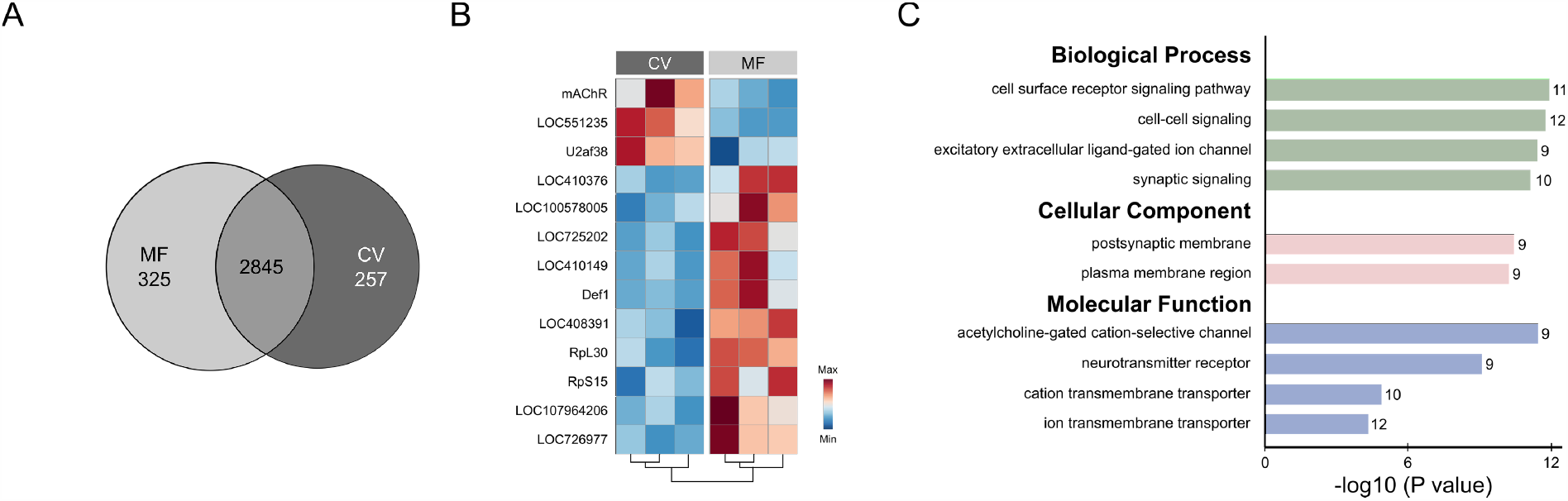
Proteomic profiling revealed upregulated neurotransmission functions in conventionalized bee brains. (**a**) Venn diagrams indicating the numbers of common and unique proteins identified in the brains of CV and MF bees. (**b**) Heatmap of differentially expressed proteins in the brains of CV and MF bees. (**c**) The most significantly enriched GO terms of identified proteins unique to the CV group (p < 0.0001, one-way ANOVA test). The number of genes involved in each GO term is displayed on the bars.

### Circulating metabolomic profiles

We have shown that gene expression, splicing, and neuronal function in the brain are influenced by the gut community, and these can be regulated by small metabolites in the circulatory system^6^. Therefore, we performed quasi-targeted metabolomics analysis of hemolymph samples from gnotobiotic bees. In total, 326 metabolites were identified among bee groups (Supplementary Data 4), and generally, the metabolic signatures of hemolymph samples were significantly different between groups (Fig. 4a, b). Interestingly, GABA and acetylcholine together with several amino acids are the most elevated metabolites in CV hemolymph (Fig. 4a). Lower levels of 5-HT are found in the hemolymph of bees colonized with *Gilliamella* and the gram-positive gut members, which is consistent with our findings of the effects on the brain neurotransmitter (Fig. 1d). To associate clusters of highly correlated metabolites to particular gut members, we performed the weighted correlation network analysis (WGCNA) based on the interaction patterns among metabolites, and bees inoculated with different gut microbes were used as the sample trait. WGCNA clustered the 326 metabolites into eight modules (M) (Supplementary Data 5), in which six modules were significantly correlated with at least one bee group (p < 0.01; Fig. 4c). The top two modules, turquoise M and blue M were both significantly associated with the CV group (Fig. 4d, Supplementary Fig. 4e), and accordingly, these two modules showed significant correlations between the metabolite significance and the intra-module connectivity for CV bees (Supplementary Fig. 4a, b). The major driving metabolites from the turquoise M and blue M are involved in the metabolism pathways of amino acid, glycerophospholipid, and carbohydrate (Fig. 4d, Supplementary Fig. 4a, b and e). The black M enriched in amino acid metabolism and some other compounds were the most correlated module to the F4 and F5 groups (Fig. 4c, d). Moreover, the Gi group was significantly associated with the red M and pink M, where more metabolites belong to the carbohydrate metabolism pathways (Fig. 4d, Supplementary Fig. 4c-e). This is consistent with the potential of *G. apicola* for carbohydrate metabolism in the gut^16,43^.

**Fig. 4.**
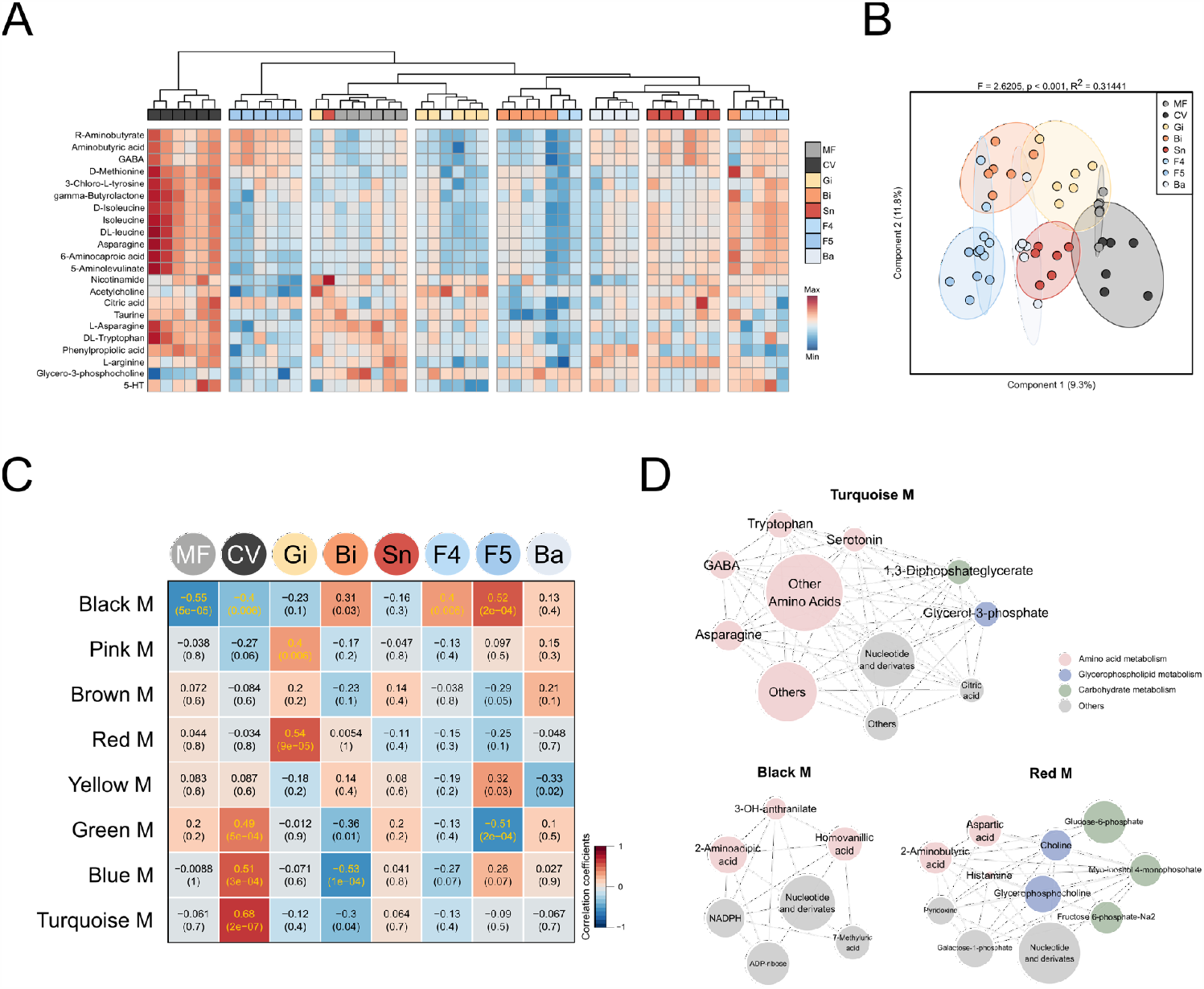
Hemolymph metabolome influenced by different honeybee gut community members. (**a**) Unsupervised hierarchical clustering heatmap of the 22 metabolites that contribute most to the separation of different groups in hemolymph samples. (**b**) Sparse PLS-DA based on all metabolites detected in the hemolymph of bees. Group differences were tested by PERMANOVA. (**c**) Weighted correlation network analysis identified eight modules (M) of metabolites highly correlated to different bee groups. Heatmap colors indicate the positive/negative Spearman’s correlation coefficient. The correlation coefficients and p values are both shown within the squares (yellow font indicates p < 0.01). (**d**) Network diagrams of differential metabolites in the turquoise, black, and red modules that are significantly correlated to CV, F4, F5, and Gi groups. Circle colors indicate different classes of metabolites in each module, and the size is proportional to the total abundance of the metabolites in the modules.

Given the evidence that the altered metabolites were largely involved in the metabolic network of neurotransmitters, we analyzed the differential level of metabolites focusing on the neurotransmitter metabolic process. Tryptophan (Trp) metabolism mainly controlled by microbiota follows three major pathways in the gastrointestinal tract: the kynurenine (Kyn) pathway, the 5-HT production pathway, and the transformation of Trp into indole by gut microbiota^44^. It showed that 3-indoleacrylic acid (IA), a key component for intestinal homeostasis^45^, was significantly elevated in the hemolymph of CV and F5 groups. While KA was not affected, Kyn was reduced in CV bees (Fig. 5). The glutamate metabolism pathway was mostly regulated in CV, Bi, Sn, and F5 groups (Fig. 5). Although glutamine was only increased in CV bees, GABA was upregulated in the hemolymph of CV, Sn, and F5 bees, which agrees with the elevated level of GABA in the brains (Fig. 1f). In addition, glycerophospholipid metabolism also plays an important role in maintaining positive mental health^46^. The hemolymph metabolites in glycerophospholipid metabolism were regulated in the CV, Gi, Sn, F5, and Ba groups compared to the MF group (Fig. 5). Specifically, acetylcholine associated with the olfactory learning and memory ability in honeybee was upregulated in the presence of *Gilliamella* (Fig. 5), coinciding with the increased level of mAChR in the brain (Fig. 3b). These results indicate that neurotransmitter related metabolisms are regulated by distinctive gut members, which may be a key mechanism of gut microbiota in modulating the brain functions.

**Fig. 5.**
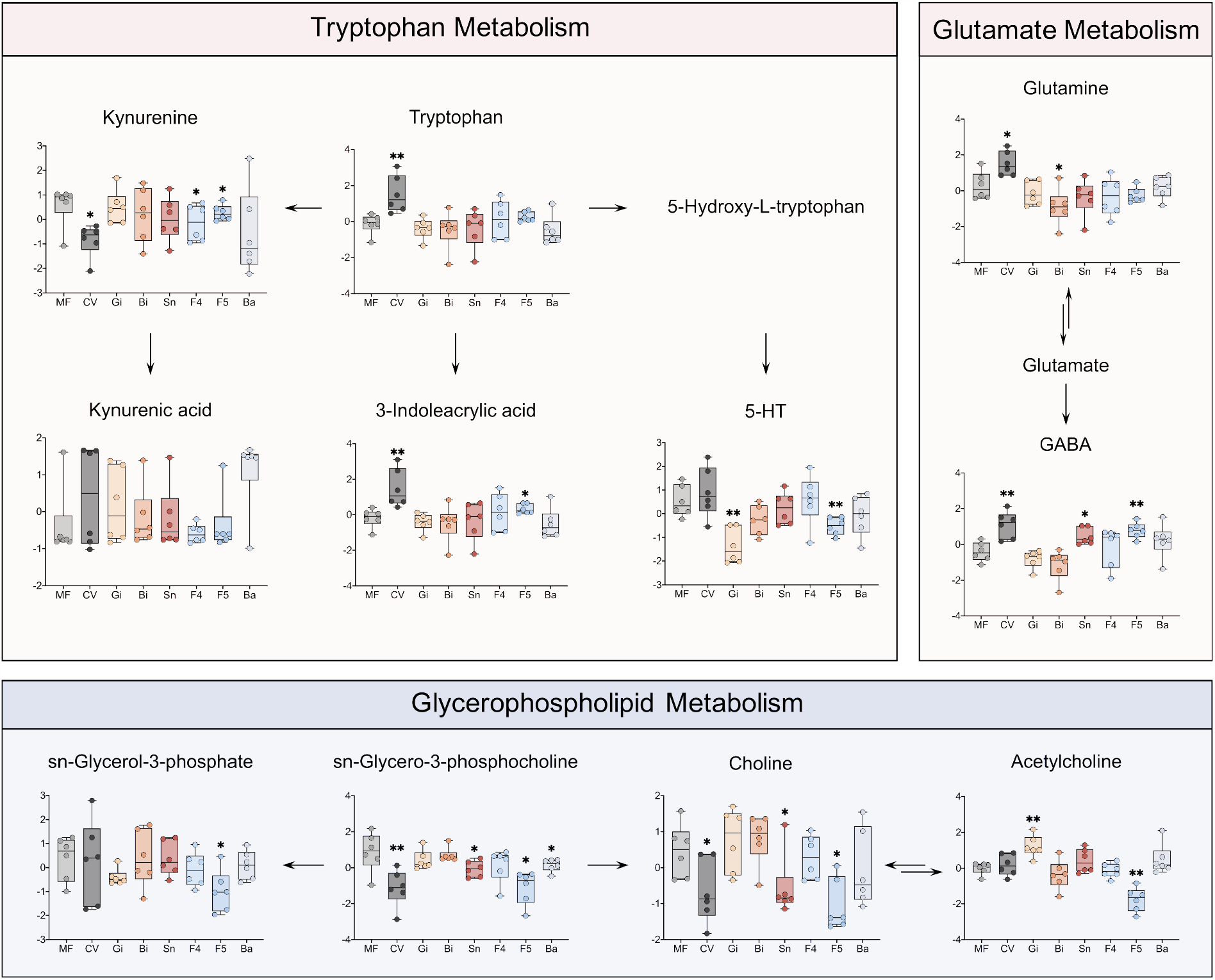
Disturbance of amino acid and glycerophospholipid metabolism pathways in honeybees colonized with different gut community members. Each dot represents the normalized concentration of differential hemolymph metabolite mapped into the tryptophan, glutamate, and glycerophospholipid metabolism pathways. Differences between MF and the other groups were tested by Mann-Whitney *u* test (*p < 0.1, **p < 0.01). Error bars represent min and max.

### Antibiotic treatment disturbs social behavior via regulating brain transcription and amino acid metabolisms

So far, our results showed that the colonization of different gut bacteria affects the honeybee behaviors under lab conditions, which is associated with the altered brain gene expression profiles, as well as the circulating and brain metabolism. We next wondered whether the perturbation of gut microbiota disturbs bee behaviors under field condition. Newly emerged bees were labeled with color tags and were then introduced to new hives with laying queens. After being returned to the hives for one week, colonies were fed wild honey or tetracycline suspended in wild honey for 5 days (Fig. 6a). The number of capped brood cells were counted, and post-treatment survival was assessed by counting the number of remaining marked bees. Although there is an increasing number of capped brood cells in the control hives, not a single capped brood was observed in the treatment group on Day 17, 18, and 19 (Fig. 6c). However, the number of recovered bees was not significantly different between control and treatment groups either before (Day 6) or after (Day 13 and 19) antibiotic treatment (Fig. 6d). Further, developing eggs and larvae were present in the brood cells of control hives with royal jelly replenished in the bottom, while only few eggs were observed in the treatment group without hatching during the whole experiment period (Fig. 6b). All these results indicate that antibiotic treatment did not obviously decrease the total number of adult bees in the hives, but disrupt the behaviors of bees without brood care. Moreover, antibiotic treatment impacts the gut appearance that the rectums of control groups are full of yellow pollens, while those of treated nursing bees were more translucent, suggesting a malnutrition status (Fig. 6b). We further characterized the composition of the gut community at both phylotype- and SDP-level through metagenomic sequencing. Although the gut community composition displayed no significant difference at pre-treatment sampling points, they displayed changes in treated bees and recovery for 7 days (Fig. 6e, Supplementary Fig. 5a). Specifically, the treatment group had a higher fraction of *Gilliamella*, while the relative abundance of *Bifidobacterium* was reduced. Tetracycline treatment affected the SDP-level profiles, and the relative abundances of Bifido-1.1, Bifido-1.2, Firm5-2, and Firm5-3 were reduced in antibiotic-treated samples (Supplementary Fig. 5b-e).

**Fig. 6.**
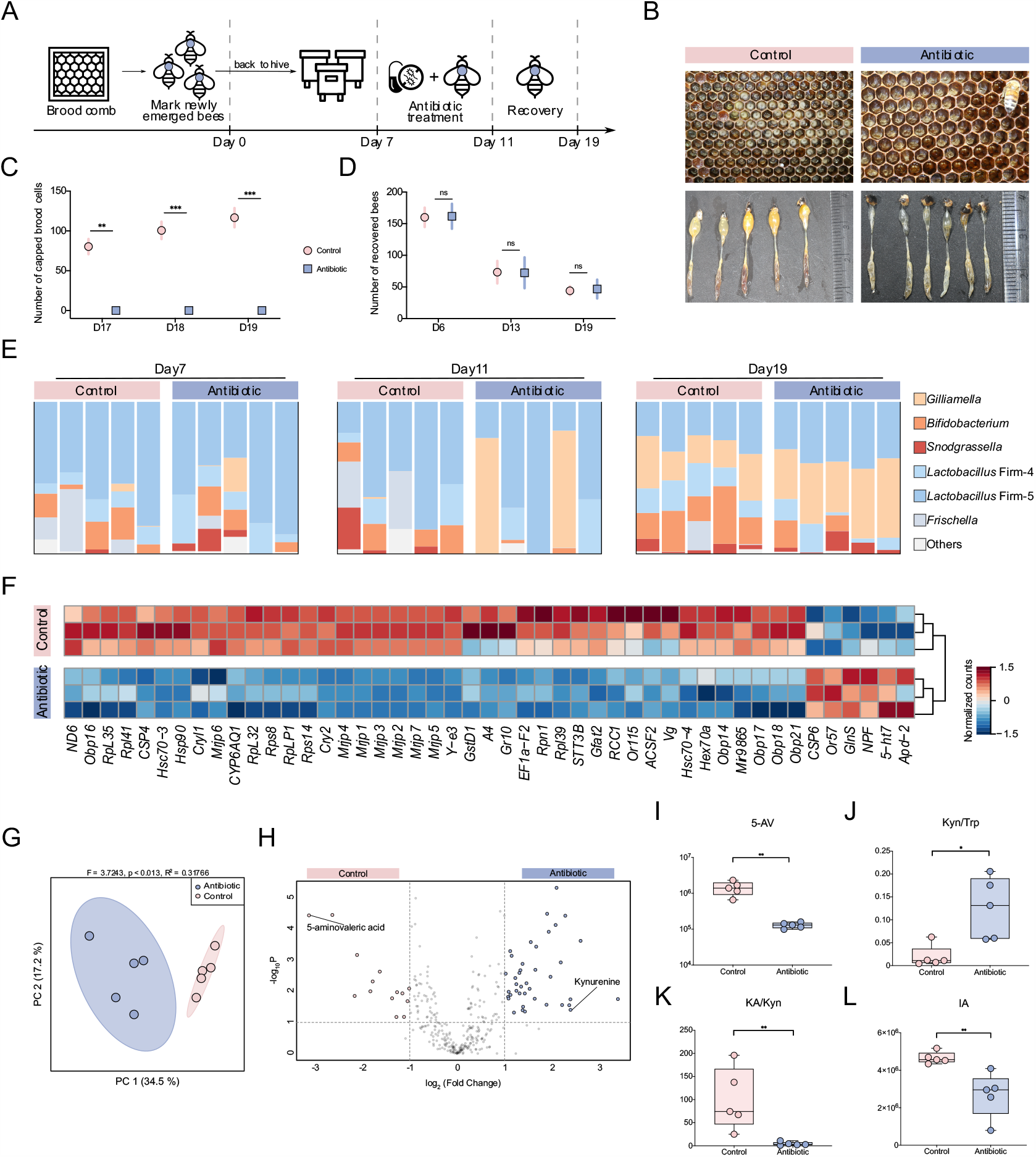
Antibiotic treatment affects honeybee social behaviors via regulating brain transcription and amino acid metabolisms. (**a**) Schematic of field experiments design for honeybee behavior. Age-controlled bees were treated with tetracycline for 5 days (Day 7– 11) in the hive and recovered for 7 days (Day 11–19). (**b**) Images of brood frames and dissected guts of control and antibiotic-treated groups. (**c**) Number of capped brood cells during the recovery stage (Day 17, 18, and 19) in three independent colonies of control and antibiotic-treated group, respectively. (**d**) The number of labelled workers recovered from the hive on Day 6 and 13 in three colonies of each group. Differences between antibiotic-treated bees and the control group were tested by multiple two-tailed t test with Benjamini-Hochberg correction (*FDR < 0.05, **FDR < 0.01, ***FDR < 0.001). (**e**) Relative abundance of phylotypes in metagenomic samples from control and antibiotic-treated groups before the antibiotic treatment (Day 7), at post-treatment (Day 11), and one week after the recovery (Day 19). (**f**) Heatmap of differentially expressed genes in the brains between control and antibiotic-treated bees. (**g**) Principal coordinate analysis based on all metabolites detected in guts of control and antibiotic-treated bees. Group differences were tested by PERMANOVA. (**h**) Volcano plot showing the differentially regulated metabolites. Metabolites significantly enriched in control bees are shown in pink, and those enriched in antibiotic-treated bees are in blue. (**i-l**) Boxplots of (**i**) the kynurenine (Kyn)/tryptophan (Trp) ratio, (**j**) the kynurenic acid (KA)/kynurenine (Kyn) ratio, the concentration of (**k**) 3-indoleacrylic acid (IA), and (**l**) 5-aminovaleric acid (5-AV) in the guts of control and antibiotic-treated bees. Group differences were tested by Mann-Whitney *u* test (*p < 0.1, **p < 0.01). Data are shown as mean ± SEM (**c-d**). Error bars represent min and max (**i-l**).

Consistent with the findings for gnotobiotic bees, we also identified a differential profile of brain gene expression for antibiotic-treated bees. The most altered genes belong to the MRJP, vitellogenin, odorant binding protein, and olfactory receptor families (Fig. 6f). Remarkably, *Gr10* and *ACSF2* that are both primarily associated with the nursing and brood-care behavior were significantly reduced by the treatment^47,48^. Additionally, we detected the differentially abundant metabolites in the gut contents of antibiotic-treated bees from the field experiment. The metabolomic profiles are clearly distinct between the two groups, and 54 metabolites were significantly different in the gut of treated bees (Fig. 6g, h). Intriguingly, 5-AV, a GABA_A_ receptor agonist affecting inhibitory GABA signaling^49^, is the most elevated compound in control bees (Fig. 6h, i). Conversely, Kyn is enriched in the gut of antibiotic-treated bees (Fig. 6h). The Kyn/Trp ratio was increased in the treatment group, while the ratio of KA/Kyn and the level of IA were decreased (Fig. 6i-k), which confirms the roles of gut microbiota in the tryptophan metabolism shift (Fig. 5). These results confirm our hypothesis that the gut microbiota affects honeybee behaviors under field conditions via the regulation of gut metabolism and gene expression in the brain.

## Discussion

Honeybees are eusocial insects that exhibit complex social, communication, and navigational behaviors with rich cognitive repertoire, such as color vision, pattern recognition, as well as learning and memory^50^. Within the colony, honeybees are characterized by the division of labor, showing striking behavioral and physiological differences between castes^51^. Although the gut microbiota composition is mostly conserved in worker bees, it differs in individuals with different behavior and physiology, such as caste, age, and worker task^52,53^, which suggests that the gut microbiota might be involved in the behavior of honeybees. While our previous study shows that bee gut microbiota alters the olfactory sensitivity^14^, the impact of microbiota on more behavioral symptoms has not been described. We report herein that a conventional gut microbiota is required for the learning ability and the establishment of memory. Although the olfactory associative learning of honeybees is largely dependent on the microbiota, the effect of gut bacteria on modulating bumblebee’s visual learning and memory is not clear^54^, suggesting that the mechanism underpinning the gut-brain interactions differs for social bees, or for olfactory and visual processing. Notably, we performed the associative appetitive learning assay in this study, while the aversive learning is also pivotal for bees to escape and avoid predators and pesticides^55^. Appetitive and aversive olfactory learning are mediated by relatively independent neural systems. Dopamine is crucial for aversive learning^56^, which is also regulated by the gut microbiota (Fig. 1e). Further evaluation of microbiota on different behaviors would assist to fully understand the mechanism of gut-brain interaction.

Olfactory and the ability of learning and memory is crucial for honeybees to cope with individual and social tasks, such as feeding and foraging^57^. Our hive experiments demonstrate that perturbation of the gut microbiota disturbs the nursing behaviors and no capped brood was observed in antibiotic-treated hives, suggesting a significant role of the normal gut microbiota in honeybee behaviors within the colony. The number of capped brood cells is a measure of the colony strength, which could also be influenced by the status of the egg-laying queen and the colony population size^58^. However, the total number of individual bees was not obviously reduced, and newly laid eggs were continuously observed in the treatment hives, implying that the perturbation of gut microbiota affects the normal hive behaviors of nurse bees and the colony reproduction.

By generating single bacterial associations, we intended to dissect the individual and combined effects of each core gut member in the sugar sensitivity. However, it showed that only conventionalized bees had a higher sensitivity, and individual gut members were not sufficient to improve the PER score, suggesting an integrative effect of the gut members, which is also reported for the *Drosophila* microbiota on the host learning^59^. Under field conditions, antibiotic treatment did not completely eliminate any core gut species but perturbed the relative abundance of the SDPs of *Lactobacillus* Firm-5 and *Bifidobacterium*, which indicates that the normal microbial community structure is required for the colony health. It has been shown that antibiotic exposure impacts bee health and dramatically reduced the survival rate with dysbiosis on the relative abundance of different bacterial genera and the fine-scale genetic diversity of the gut community^60^. In addition to the increased susceptibility to ubiquitous opportunistic pathogens^61^, the colony losses resulting from antibiotic treatment could be partly due to the altered hive behaviors.

Our RNA-seq analysis of gnotobiotic bee brains showed that numerous transcripts differed in expression levels, moreover, genes related to honeybee labor division and olfactory ability were altered by gut bacteria. For example, genes encoding the MRJPs involved in the learning and memory abilities of honeybees were upregulated in bacteria-colonized bees. Consistently, it has been reported that the expression level of *mrjp1* and *mrjp4* are repressed in the brains of imidacloprid treated bees, which also exhibit impaired learning^27^. The expression level of *Vg* was disturbed by gut microbiota in both laboratory experiments and in the hive, corroborating with the previous finding in MF bee abdomen^14^. Vitellogenin is a nutritional status regulator that influences honeybee social organization and stress resilience^62^. The expression of *Vg* and survivorship could be elevated by the addition of pollen to the diet^63^, whereas the effect is alleviated by the disturbance of gut community^64^, indicating the important role of gut microbiota in *Vg* regulation via the nutritional metabolism.

Homologous molecular mechanism in social responsiveness has been documented between honeybee and human^34^. The transcription profile in brains of bees with disordered social behaviors is distinct from that of normal bees, and differently expressed genes in unresponsive individuals are enriched for human ASD-related genes. These genes are also found associated with the polymorphism of the halictid bee *Lasioglossum albipes*, indicating their implications in social behaviors^65^. Despite the disturbed gene expression level, AS patterns of the ASD-related genes are also highly correlated to mental disorders^7^. In our dataset, the analysis of gene splicing identified extensive differences among bacteria-colonized groups of bees, and the altered genes compared to MF bees largely overlapped with the SPARK and SFARI gene datasets for Autism (Fig. 2). The MXE and SE events of two high-confidence ASD risk genes, *Ank2* and *Rims1*, were predominantly affected by the gut bacteria. These two genes are also found affected in the brains of mice colonized by ASD-human gut microbiota^7^. Correspondingly, our brain proteomics revealed that the splicing factor was upregulated in CV brains, supporting the contribution of microbiota to splicing regulation. All these findings demonstrate a deep conservation for genes related to social responsiveness of human and distantly related insect species, and reflect a common role of the gut microbes implicated in the evolution of sociality^66^.

Neurotransmitters that carry and pass information between neurons are essential for brain functions, which are important modulators of behaviors. In honeybees, four monoamine neurotransmitters play important roles in learning and memory^22^, and olfactory sensitivity^22,23^. In addition, GABA and acetylcholine have been physiologically characterized to induce currents between neurons within the olfactory pathways and contribute to the odor memory formation^42^. Concentrations of most identified neurotransmitters were regulated by different gut members, corroborating with the roles of gut microbiota in the altered behaviors in the lab and hive experiments. Alternatively, it recently shows that nestmate recognition cues are defined by gut bacteria, possibly by modulating the host metabolism or by the direct generation of the colony-specific blends of cuticular hydrocarbon^67^. In leaf-cutting ants and termites, gut microbiota suppression by antibiotics also influences the recognition behavior toward nestmates, which may be directed by the bacterial metabolites as recognition cues in the feces^68,69^. Nevertheless, the effect of gut community is mainly driven by the microbial metabolism, specifically the amino acid and lipid metabolic pathways, which can further influence the circulation system and the synthesis of neuroactive molecules of the host. Perturbation in gut tryptophan metabolism has been associated with neuropsychiatric disorders in human and *Drosophila* model, characterized by reduced plasma level of tryptophan^70^, high IDO1 activity^71^, and high level of 5-HT in brains^72^. In honeybees, 5-HT was also elevated in MF bee brains compared to bacteria-colonized counterparts, moreover, the level of IDO1 activity (assessed by Kyn/Trp ratio) was higher in the gut of antibiotic-treated bees in the field colony. In addition, acetylcholine synthesized in the glycerophospholipid pathway is a neurotransmitter crucial for the olfactory learning and memory ability in honeybees. Our results revealed that gut microbiota mediated the cholinergic metabolism in the hemolymph, and correspondingly, the brain proteomics showed an increased level of the muscarinic acetylcholine receptor in CV bees. Cholinergic signaling via the mAChR is critical for the olfactory associative learning and foraging behaviors^41^. Moreover, the stimulation of the mAChR of honeybee increases the volume of the mushroom body neuropil, which mimics the reinforcement of cholinergic neurotransmission in foraging bees^73^. A reduced mushroom body calycal growth is also associated with lower learning performance in bumblebees through micro-computed tomography scanning^74^. It would be interesting to investigate whether gut microbes impact the structural changes of the brain in future studies.

It is increasingly realized that gut microorganisms may influence the development of social behaviors across diverse animal hosts^66^. While hypothesis-generating, translating these correlations into actionable outcomes is challenging in humans. Honeybees are colonial and highly social with multiple symbolic behaviors, which offer an experimental tool to investigate the relationship between the microbiota and host brain functions and help to uncover the causal mechanisms underlying sociability. Our study highlights multiple parallels between honeybee and human that gut microbiota plays an important role in host brain functions. The development of genetic tools manipulating both the bee host and the gut bacteria would facilitate the investigation of the molecular basis of host-microbe interactions via the gut-brain axis^75,76^.

## Methods

### Generation of microbiota-free, mono-colonized and conventionalized honeybees

Microbiota-free (MF) bees were obtained as described by Zheng *et al*.^16^ with modifications. Late-stage pupae were removed manually from brood frames and placed in sterile plastic bins. The pupae emerged in an incubator at 35°C, with humidity of 50%. Newly emerged MF bees (Day 0) were kept in axenic cup cages with sterilized sucrose syrup (50%, wt/vol) for 24 h and divided into three groups: 1) MF, 2) mono-colonized (MC) and 3) conventional (CV) bees. For each setup, 20–25 MF bees (Day 1) were placed into one cup cage, and the bees were feeding on the corresponding solutions or suspensions for 24 h. For the MF group, 1 mL of 1×PBS was mixed with 1 mL of sterilized sucrose solution (50%, wt/vol) and 0.3 g sterilized pollen. For the MC group, stocks of *Gilliamella apicola* (W8127), *Snodgrassella alvi* (W6238G3), *Bifidoobacterium asteroides* (W8113), *Bartonella apis* (B10834G6), *Lactobacillus* sp. Firm-4 (W8089), and *Lactobacillus* sp. Firm-5 (W8172) in 25% glycerol stock at –80°C were resuspended in 1mL 1×PBS (Solarbio, Beijing, China) at a final OD_600nm_ of 1, and then mixed with 1 mL sterilized sucrose solution (50%, wt/vol) and 0.3 g sterilized pollen. For the CV group, 5 µL homogenates of freshly dissected hindguts of nurse bees from their hives of origin were mixed with 1 mL 1×PBS, 1 mL sterilized sucrose solution (50%, wt/vol) and 0.3 g sterilized pollen. Then MF, MC, and CV bees were provided sterilized sucrose (0.5 M) with sterile pollens and kept in an incubator (35°C, RH 50%) until day 7. Brains, guts, and hemolymph of bees were collected on day 7 for further analysis.

### Bacterial load quantification

Colonization levels of MF and MC bees were determined by colony-forming units from dissected guts, as described by Kwong *et al*.^77^. Colonization levels of CV bees were determined by quantitative PCR as previously described by Engel *et al*.^13^. All qPCR reactions were carried out in a 96-well plate on the StepOnePlus Real-Time PCR system (Applied Biosystems; Bedford, MA, USA) with the thermal cycling conditions as follows: denaturation stage at 50°C for 2 min followed by 95°C for 2 min, 40 amplification cycles at 95°C for 15 s, and 60°C for 1 min. Melting curves were generated after each run (95°C for 15 s, 60°C for 20 s and increments of 0.3°C until reaching 95°C for 15 s) to compare dissociation characteristics of the PCR products obtained from gut samples and positive control. Each reaction was performed in triplicates on the same plate in a total volume of 10 μl (0.2 μM of each forward and reverse primer; and 1x SYBR® Select Master Mix, Applied Biosystems; Bedford, MA, USA) with 1 μl of DNA or cDNA (to assess virus loads). Each plate contained a positive control and a water control. After the calculation of the bacterial 16S rRNA gene copies, normalization with the actin gene was carried out to reduce the effect of gut size variation and extraction efficiency. In brief, bacterial 16S rRNA gene copies were normalized to the medium number of actin gene copies by dividing by the ‘raw’ copy number of actin for the given sample and multiplying by the median number of actin gene copies across all samples. Universal bacteria primers (Forward: 5’ -AGGATTAGATACCCTGGTAGTCC-3’, Reverse: 5’-YCGTACTCCCCAGGCGG-3’)^13^ and *Apis mellifera* actin (Forward: 5’ -TGCCAACACTGTCCTTTCTG -3’, Reverse: 5’-AGAATTGACCCACCAATCCA -3’)^78^ were used here.

### Tissue collection

The whole guts were dissected by tweezers disinfected with 75% alcohol. Dissected guts were directly crushed in 25% (vol/vol) glycerol on ice for bacterial load quantification or collected into an empty 1.5-mL centrifuge tube for metagenomic sequencing and metabolomics analysis. All gut samples were frozen at –80°C until analysis. Honeybee brains were collected using a dissecting microscope (Canon). Individual bee was fixed on beeswax using two insect needles through the thorax. After removing the head cuticle, the whole brain was placed on a glass slide and soaked in RNAlater (Thermo; Waltham, MA, USA) or proteinase inhibitor (Roche; Mannheim, Germany) for gene expression profiling, proteome analysis, and neurotransmitters concentration quantification. Then hypopharygeal glands, salivary glands, three simple eyes, and two compound eyes were carefully removed. Dissected brains were kept frozen in −80°C. Hemolymph was collected using a 10 μL pipettor (Eppendorf; Hamburg, Germany) from the incision above the median ocellus. A minimum of 50 μL of hemolymph was collected from10 bees into a 1.5-mL centrifuge tube. During the collection process, tubes are temporarily preserved on dry ice and subsequently stored at −80 °C until analysis.

### I n laboratory honeybee behavior experiment

#### Learning and memory

We measured the olfactory learning and memory ability of seven-day-old MF, CV, and CV+tet bees. MF and CV bees were generated as described above. CV+tet bees were fed 450 μg/ml (final concentration) of tetracycline suspended in sterilized 0.5 M sucrose syrup on Day 5 after the eclosion for 24 h and then were fed sucrose syrup for another 24 h for recovery. Experiments of olfactory learning and memory were performed as previously described^20,79^ with modifications (Fig. 1a). In brief, bees were starved for 2 h by removing sugar syrup and bee bread from the cup cage before the test and were then mounted to modified 0.8 mm wide bullet shell with sticky tape restraining harnesses (Supplementary Movie 1). The whole experiment was performed in a room with a stable light source at room temperature. Each bee individual was checked for their intact proboscis extension response by touching the antennae with 50% sucrose solution without subsequent feeding 15 min before the experiment. Nonanol (olfactory learning; Sigma-Aldrich; Saint Louis, MO, USA) and hexanal (negative control; Macklin; Shanghai, China), which could be distinguished by honeybee^80^, were used as odor sources. The odor was produced by pricking holes on a 0.8 cm wide filter paper and soaking it in 0.5 mL nonanol or hexanal, and the filter paper was then slipped into a 10 mL injector. During conditioning, individual harnessed bee was placed in front of an exhaust fan to prevent odor build-up in subsequent experiments. Bees were trained for 10 trials with an inter-trial interval of 10 min to associate nonanol odor as conditioned stimulus with a reward of 50% sucrose solution as unconditioned stimulus.

At the beginning of each trial, the harnessed bee was placed inside the arena for 5 sec to allow familiarization with the experimental context. Thereafter, the nonanol odor was presented before its antennal for 6 sec, and then 0.4 uL droplet of sucrose solution was delivered to the bee using a syringe needle, which directly touched the proboscis to evoke PER. Once the 10 trials of a conditioning session were completed, bees were kept in the dark without being fed for 3 h. Two unreinforced olfactory memory tests were administered 3 h after olfactory conditioning: one with the conditioned stimulus odor (nonanol) and one with a novel odor (hexanal). The order of presentation was randomized across subjects. A clean and tasteless injector was delivered to the bee after each odor test to exclude visual memory of reward during olfactory conditioning. Bees only extending the proboscis to nonanol odor were considered as successful memorized individuals (Fig. 1a).

#### Gustatory responsiveness

Seven-day-old MF, MC, and CV bees were used to measure the response to different concentrations of sucrose solution as previously described with some modifications^14^. Before the test, bees were starved for 2h in the incubator by removing sugar syrup and bee bread from the cup cage. Bees were then mounted to modified 2.0-mL centrifuge tubes using Parafilm M (Bemis; Sheboygan Falls, WI, USA), and they could only move their heads and propodeum for antennae sanitation. Individual responsiveness was measured by presenting a series concentration of sucrose solutions (0, 0.1, 0.3, 1, 3, 10, and 30%; wt/vol) to the antennae of bees^81^. Before each sucrose solution presentation, all bees were tested for their response to pure water in order to control for the potential effects of repeated sucrose stimulations that may lead to either sensitization or habituation^82^. The inter-stimulus interval between water and sucrose solution was 4 min. When a bee’s antenna is stimulated with a sucrose solution of sufficient concentration, the bee reflexively extends its proboscis. The lowest sucrose concentration at which an individual responded by extending its proboscis was recorded and interpreted as its sugar response threshold. At the end of the experiment, a gustatory response score was obtained for each bee, which is based on the number of sucrose concentrations to which the bees responded. The response was arbitrarily quantified with scores from 1 to 7, where 1 represented a bee that only responded to the highest sucrose concentration, while a score of 7 represented an individual that responded to all concentrations tested. If a bee failed to respond in the middle of a response series, this ‘failed’ response was considered to be an error and the bee was deemed to have responded to that concentration as well. Bees that did not respond to any of the sucrose concentrations were excluded from further analyses. In addition, bees that responded to all concentrations of sucrose solutions and all presentations of water were also excluded as they appeared not to be able to discriminate between sucrose and water^82^.

#### Hive behavior experiment

The fieldwork took place in 2019 at the apiary of China Agricultural University, Beijing, China, and the experiment was performed twice in July and August, respectively. To observe the effect of gut microbiota on the hive bee behaviors with the same age, two independent single-cohort colonies were set up as previously described^83^. Briefly, brood frames were collected from a single hive and adult bees were brushed off. The frames were then kept in the laboratory incubating at 35°C and 50% relative humidity. In two days, about 1,000 bees emerged from each frame in the incubator, and we labeled 300 individuals with colored tags on their thorax. All newly emerged bees were then introduced to new empty hives together with a newly mated laying queen^84^. Two hives for control and treatment were established. Control colony bees were fed wild honey along with the whole experiment, and treatment groups were fed wild honey suspended with 450 ug/ml of tetracycline (final concentration) from Day 7 after the establishment of hives (Fig. 6a), and the antibiotic treatment lasted for 5 days. The number of capped brood cells was counted every day, and post-treatment survival in the hive was assessed by counting the number of remaining marked bees of the whole hive^61^. Marked bees for both control and treatment groups were collected from each hive at time points of 7, 11, and 19 day following the set-up of hives, and the hind guts and brain tissue were dissected. All samples were stored at −80 °C until analysis.

#### Gut DNA extraction and metagenomic sequencing

Bee individuals of either control or antibiotic groups were sampled on day 7, 11, and 19 during the hive behavior experiment (Fig. 6a). Total genomic DNA of the gut microbiota was extracted from the whole gut homogenate using CTAB method as previously described^14^. DNA samples were sent to Novogene Bioinformatics Technology Co. Ltd. (Beijing, China) for shotgun metagenome sequencing. Sequencing libraries were generated using NEBNext Ultra™ II DNA Library Prep Kit for Illumina (New England Biolabs; Ipswich, MA, USA), and the library quality was assessed on Qubit 3.0 Fluorometer (Life Technologies; Grand Island, NY, USA) and Agilent 4200 (Agilent, Santa Clara, CA) system. The libraries were then sequenced on the Illumina Novaseq 6000 platform (Illumina; San Diego, CA, USA) and 150 bp paired-end reads were generated. The SDP- and phylotype-level community structure of each metagenomic sample was profiled following the Metagenomic Intra-Species Diversity Analysis System (MIDAS) pipeline^85^. A custom bee gut bacteria genomic database was generated based on 407 bacterial isolates from honeybees and bumblebees (Supplementary Data 6). Before the classification, we removed reads belonging to the honeybee reference genome (version Amel_HAv3.1) using KneadData v 0.7.3. We then ran the ‘species’ module of the ‘run_midas.py’ and ‘merge_midas.py’ scripts in MIDAS with our custom bacterial genome database, which aligned reads to universal single-copy gene families of phylogenetic marker genes using HS-BLASTN to estimate the abundance of phylotypes and SDPs for each sample. Local alignments that cover < 70% of the read or fail to satisfy the gene-specific species-level percent identity cut-offs were discarded.

#### Brain gene expression analysis

Total RNA was extracted from individual brains using the Quick-RNA MiniPrep kit (Zymo; Irvine, CA, USA). RNA degradation and contamination were monitored on 1% agarose gels, and the purity was checked with the NanoPhotometer spectrophotometer (IMPLEN; CA, USA). RNA integrity was assessed using the RNA Nano 6000 Assay Kit of the Bioanalyzer 2100 system (Agilent Technologies; Santa Clara, CA, USA). RNA sequencing libraries were generated using NEBNext Ultra RNA Library Prep Kit for Illumina (New England BioLabs; Ipswich, MA, USA) and index codes were added to attribute sequences to each sample. The clustering of the index-coded samples was performed on a cBot Cluster Generation System using TruSeq PE Cluster Kit v3-cBot-HS (Illumina; San Diego, CA, USA), and the library preparations were then sequenced on an Illumina NovaSeq 6000 platform (Illumina; San Diego, CA, USA) and 150 bp paired-end reads were generated. Sequencing quality of individual samples was assessed using FastQC v0.11.5 with default parameters. An index of the bee reference genome (Amel_HAv3.1) was built using HISAT2 v2.0.5^86^, and the FastQC trimmed reads were then aligned to the built index using HISAT2 v2.1.0 with default parameters. Gene expression was quantified using HTSeq v0.7.2^87^ with mode ‘union’, only reads mapping unambiguously to a single gene are counted, whereas reads aligned to multiple positions or overlapping with more than one gene are discarded. If it were counted for both genes, the extra reads from the differentially expressed gene may cause the other gene to be wrongly called differentially expressed, so we chose ‘union’ mode.

Differential gene expression analysis was performed using the DESeq2 package^88^ in R. We modeled read counts following a negative binomial distribution with normalized counts and dispersion. The proportion of the gene counts in the sample to the concentration of cDNA was scaled by a normalization factor using the median-of-ratios method. The variability between replicates is modeled by the dispersion parameter using empirical Bayes shrinkage estimation. For each gene, we fit a generalized linear model to get the overall expression strength of the gene and the log 2-fold change between CV, MC, and MF groups. For significance testing, differential gene expression is determined by the Wald test. The resulting p-values were corrected for multiple comparisons using the Benjamini-Hochberg FDR method^89^. Genes with an adjusted P-value < 0.05 and |log_2_FoldChange| > 1 were assigned as differentially expressed.

To get a better annotation of the honeybee reference genome, we re-annotate it using eggNOG-mapper v5.0^90^. 6,269 out of 12,375 honeybee genes were successfully assigned to a KO entry with the ‘diamond’ mode, and the hierarchy information of the KEGG metabolic pathway was extracted. Functional analysis of differentially expressed genes was performed based on KEGG Orthologue (KO) markers. The percentages of KO markers belong to each category (KEGG Class at level 3) out of total MC-, CV-, and MF-enriched KO markers were designated as a comparison parameter. The significance level was calculated by Fisher’s exact test using clusterProfiler v3.10.1^91^.

Analysis of event-level differential splicing was performed using rMATS v4.0.2^92^ based on the bee reference genome. An exon-based ratio metric, commonly defined as percent-spliced-in value, was employed to measure the alternative splicing events. The percent spliced in (PSI) value is calculated as follows:

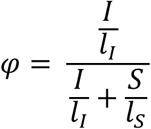

, where S and I are the numbers of reads mapped to the junction supporting skipping and inclusion form, respectively. Effective length l is used for normalization. The PSI value was calculated for several classes of alternative splicing events, including skipped exon (SE), alternative 5’ splice site (A5SS), alternative 3’ splice site (A3SS), mutually exclusive exons (MXE), and retained introns (RI). Events with p < 0.05 were considered differentially spliced across gnotobiotic bees and microbiota-free bees.

To find the overlaps between the differentially expressed or spliced genes of bee brain and those from humans autism spectrum disorders, a total of 3,531 high-quality reference protein sequences corresponding to 948 known autism risk genes (SFARI: https://gene.sfari.org/, SPARK for Autism: http://spark-sf.s3.amazonaws.com/SPARK_gene_list.pdf) were aligned against the protein sequences of honeybee genome using BLASTP^93^ with two-way best matching strategy. In total, 649 autism protein sequences obtained a match (Similarity > 30% and e-value < 0.000394). Then we calculated the intersection of the autism risk genes and the differentially expressed or spliced genes between bacteria colonized bees and MF bees (p < 0.05).

### Brain proteome analysis

The proteome analysis was performed as described by Meng *et al*.^94^. Briefly, three replicates per treatment group were analyzed for each group of bees. 20 dissected honeybee brains were pestle ground, sonicated, and cooled on ice for 30 min in a lysis buffer (8 M urea, 2 M thiourea, 4% 3-((3-cholamidopropyl) dimethylammonio)-1-propanesulfonate acid (CHAPS), 20 mM tris-base, 30 mM dithiothreitol (DDT)). The homogenate was centrifuged at 12,000 g and 4 °C for 15 min, followed by supernatant recovery. Then 4 volumes of ice-cold acetone were added for 30 min to precipitate protein. The protein pellets were collected after centrifugation (8,000g, 4°C for 15 min), then dried at room temperature, and dissolved in 40 mM NH_4_HCO_3_. To prevent reformation of disulfide bonds, the dissolved protein samples were incubated with 100 mM of DDT (DDT/protein (V: V=1:10)) for 1 h and then alkylated with 50 mM of iodoacetamide (IAA) (DDT/IAA (V: V=1:5)) for 1 h in the dark. Finally, the resultant protein was digested with trypsin (enzyme: protein (W: W=1:50)) at 37°C for 14 h. After digestion, the enzymatic reaction was stopped by adding 1 μL of formic acid into the mixture. The digested peptides were centrifuged at 13,000g and 4°C for 10 min. The supernatant was recovered and extracted using a SpeedVac system (RVC 2-18, Marin Christ; Osterod, Germany) for subsequent LC-MS/MS analysis.

Peptides were measured by the EASY-nLC 1000 liquid chromatograph (Thermo Fisher Scientific, Waltham, MA, USA) on a Q Exactive HF mass spectrometer (Thermo Fisher Scientific). Peptides were separated on an analytical column packed with 2 μm Aqua C18 beads (15cm long, 50 μm inner diameter, Thermo Fisher Scientific) at a flow rate of 350 nL/min, using a 120-min gradient (2% (vol/vol) to 10% (vol/vol) acetonitrile with 0.1% (vol/vol) formic acid). The Q Exactive was operated in the data-dependent mode with the following settings: 70000 resolution, 350–1,600 *m/z* full scan, Top 20, and a 2 *m/z* isolation window. Identification and label-free quantification of peptides were done with PEAKS Studio X+ (Bioinformatics Solutions Inc.; Waterloo, ON, Canada) against the sequence database (21,780 protein sequences of *Apis mellifera*), coupled with a common repository of adventitious proteins database (cRAP, https://www.thegpm.org/dsotw_2012.html). The search parameters were: parent ion tolerance, 15 ppm; fragment tolerance, 0.05 Da; enzyme, trypsin; maximum missed cleavages, 3; fixed modification, carbamidomethyl (C, +57.02 Da); and variable modification, oxidation (M, +15.99 Da). A protein was confidently identified only if it contained at least one unique peptide with at least two spectra, applying a threshold of false discovery rate (FDR) ≤ 1.0% by a fusion-decoy database searching strategy (PMID: 22186715). Proteins significantly differential between groups were identified using ANOVA (p-value < 0.05 and a fold change of ≥ 1.5).

The functional gene ontology (GO) term and pathway were assessed using ClueGOv2.5.5, Cytoscape plug-in software (http://www.ici.upmc.fr/cluego/). The analysis was performed by comparing an input data set of identified proteins to all functionally annotated GO categories in the entire genome of *Apis mellifera* from UniProt. The significantly enriched GO terms in cellular component (CC), molecular function (MF), biological processes (BPs) and pathways were reported using a two-sided hyper-geometric test and only a p-value ≤ 0.05 was considered. Then, Bonferroni step-down was used to correct the p-value to control FDR. Functional grouping of the terms was based on the GO hierarchy. The tree level was ranged from 3 to 8, and the kappa score level was 0.4.

### Targeted metabolomics for brain neurotransmitters

Brain tissues dissected from MF, MC, and CV bees were sent to Biotree Biotech Co. Ltd. (Shanghai, China) for targeted metabolomics analysis of dopamine, octopamine, serotonin, tyramine, and GABA. Six brain tissues from one treatment group were put into one tube and centrifuged (2400 g × 1 min at 4 °C). 100 µL acetonitrile containing 0.1% formic acid and 20 µL ultrapure water were added and the tubes were vortexed thoroughly. Metabolites were sonicated in an ice-water bath for 30 min, followed by subsiding at −20 °C for 2 h. Supernatants were collected after centrifugation (14,000 g × 10 min at 4 °C). 20 µL of supernatant were transferred to a new vial followed by incubation for 30 min after the addition of 10 µL sodium carbonate solution (100 mM) and 10 µL 2% benzoyl chloride acetonitrile. Then 1.6 µL internal standard and 20 µL 0.1% formic acid were added, and the samples were centrifuged (14,000 g × 5 min at 4 °C). 40 μL of the supernatants were transferred to an auto-sampler vial for downstream UHPLC-MS/MS analysis. Serotonin hydrochloride, γ-aminobutyric acid, dopamine hydrochloride, tyramine, and octopamine hydrochloride (Aladdin; Shanghai, China) derivatized with benzoyl chloride (Sigma-Aldrich; Saint Louis, MO, USA) were used for the construction of the calibration standard curve. The internal standards mixture (γ-aminobutyric acid, dopamine hydrochloride, serotonin hydrochloride, tyramine, and octopamine hydrochloride derivatized with benzoyl chloride-d5 (Sigma-Aldrich; Saint Louis, MO, USA)^95^ of the corresponding concentration were prepared, respectively.

The UHPLC separation was carried out using an ExionLC System (AB SCIEX; MA, USA), and the samples were analyzed on the QTRAP 6500 LC-MS/MS system (AB Sciex; Framingham, MA, USA). 2 μL of samples were directly injected onto an ACQUITY UPLC HSS T3 column (100 × 2.1 mm × 1.8 μm; Waters; Milford, Ma, USA). The column temperature was set at 40 °C, and the auto-sampler temperature was set at 4 °C. Chromatographic separation was achieved using a 0.30 ml/min flow rate and a linear gradient of 0 to 2% B within 2 min; 2%–98% B in 9 min, followed by 98% B for 2 min and equilibration for 2 min. Solvent A is 0.1% formic acid and solvent B is acetonitrile. For all multiple reaction monitoring (MRM) experiments, 6500 QTrap acquisition parameters were as follows: 5000 V Ion-spray voltage, curtain gas setting of 35 and nebulizer gas setting of 60, temperature at 400 °C. Raw data were analyzed using Skyline^96^.

### Quasi-Targeted metabolomics analysis

Hemolymph and gut homogenate metabolites were determined by quasi-targeted metabolomics by HPLC-MS/MS. Gut samples (100mg) were individually grounded with liquid nitrogen and the homogenate was resuspended with prechilled 500 μL 80% methanol and 0.1% formic acid by well vortexing. 50 μL of hemolymph samples were mixed with 400 μL prechilled methanol by vortexing. All samples were incubated on ice for 5 min and then centrifuged at 15,000 × g, at 4°C for 10 min. The supernatant was diluted to a final concentration containing 53% methanol by LC-MS grade water. The samples were then transferred to a fresh vial and centrifuged at 15,000 × g, 4°C for 20 min. Finally, the supernatant was injected into the LC-MS/MS system, and the analyses were performed using an ExionLC AD system (SCIEX) coupled with a QTRAP 6500+ mass spectrometer (SCIEX). Samples were injected onto a BEH C8 Column (100 mm × 2.1 mm × 1.9 μm) using a 30-min linear gradient at a flow rate of 0.35 mL/min for the positive polarity mode. Eluent A was 0.1% formic acid-water and eluent B is 0.1% formic acid-acetonitrile. The solvent gradient was set as follows: 5% B, 1 min; 5-100% B, 24.0 min; 100% B, 28.0 min;100-5% B, 28.1 min;5% B, 30 min. QTRAP 6500+ mass spectrometer was operated in positive polarity mode with curtain gas of 35 psi, collision gas of Medium, ion spray voltage of 5500V, temperature of 500°C, ion source gas of 1:55, and ion source gas of 2:55. For negative ion mode, samples were injected onto aHSS T3 Column (100 mm × 2.1 mm) using a 25-min linear gradient at a flow rate of 0.35 mL/min. The solvent gradient was set as follows: 2% B, 1 min; 2%–100% B, 18.0 min; 100% B, 22.0 min; 100%–5% B, 22.1 min; 5% B, 25 min. QTRAP 6500+ mass spectrometer was operated in negative polarity mode with curtain gas of 35 psi, collision gas of medium, ion spray voltage of −4500V, temperature of 500°C, ion source gas of 1:55, and ion source gas of 2:55.

Detection of the experimental samples using MRM was based on Novogene in-house database. Q3 (daughter) was used for the quantification. Q1 (parent ion), Q3, retention time, declustering potential, and collision energy were used for metabolite identification. Data files generated by HPLC-MS/MS were processed with SCIEX OS (version 1.4) to integrate and correct the peaks. A total of 326 compounds were identified in the hemolymph samples. Metabolomics data analysis was then performed using MetaboAnalyst 4.0^97^.

### Weighted gene co-expression network analysis (WGCNA)

R software package WGCNA 1.69^98^ was used to identify key phenotype-related metabolic modules based on correlation patterns. The Pearson correlation matrix was calculated for all possible metabolite pairs and then transformed into an adjacency matrix with a soft thresholding power setting to 5 for the best topological overlap matrix. A dynamic tree cut algorithm was used to detect groups of highly correlated metabolites. The minimum module size was set to 14 and the threshold for merging module was set to 0.25 as default. Each module was assigned a unique color and contained a unique set of metabolites. After obtaining modules from each group, module eigenmetabolite was calculated with the “ModuleEigengenes” function. Association analysis between a module and the trait of each group was performed using the function of “corPvalueStudent” based on the module eigenmetabolite. p < 0.01 was set for statistical significance. Metabolites in each module were annotated on the KEGG Database and classified into major categories using MetaboAnalyst 4.0^97^ for enrichment analysis. Finally, the network connections among metabolites in modules were visualized using Cytoscape 3.7.0^99^.

### Statistical analysis

Comparison of the learning and memory results was tested by Chi-squared test using GraphPad Prism 8.2.0 software. Comparisons of the distribution of gustatory response score, neurotransmitters, normalized and raw metabolite data of different bacterial colonized groups were made by Mann–Whitney *u* test using GraphPad Prism 8.2.0 software. The exact value of n representing the number of groups in the experiments described was indicated in the figure legends. Any additional technical replicates are described within the Methods and the Results.

### Data Availability

The raw data for outdoor honeybee gut microbiome shotgun sequencing has been deposited under BioProject PRJNA670603. The accession numbers for the RNA sequencing data are PRJNA670620 and PRJNA668910. The proteomic data has been deposited to the Proteome Xchange Consortium with the dataset identifier PXD022304.

### Code availability

The list of analysis software and all scripts generated for analysis have been deposited on GitHub at: https://github.com/ZijingZhang93/bee_BGA.git.

## Supplemental Information

**Supplementary Fig. 1.**
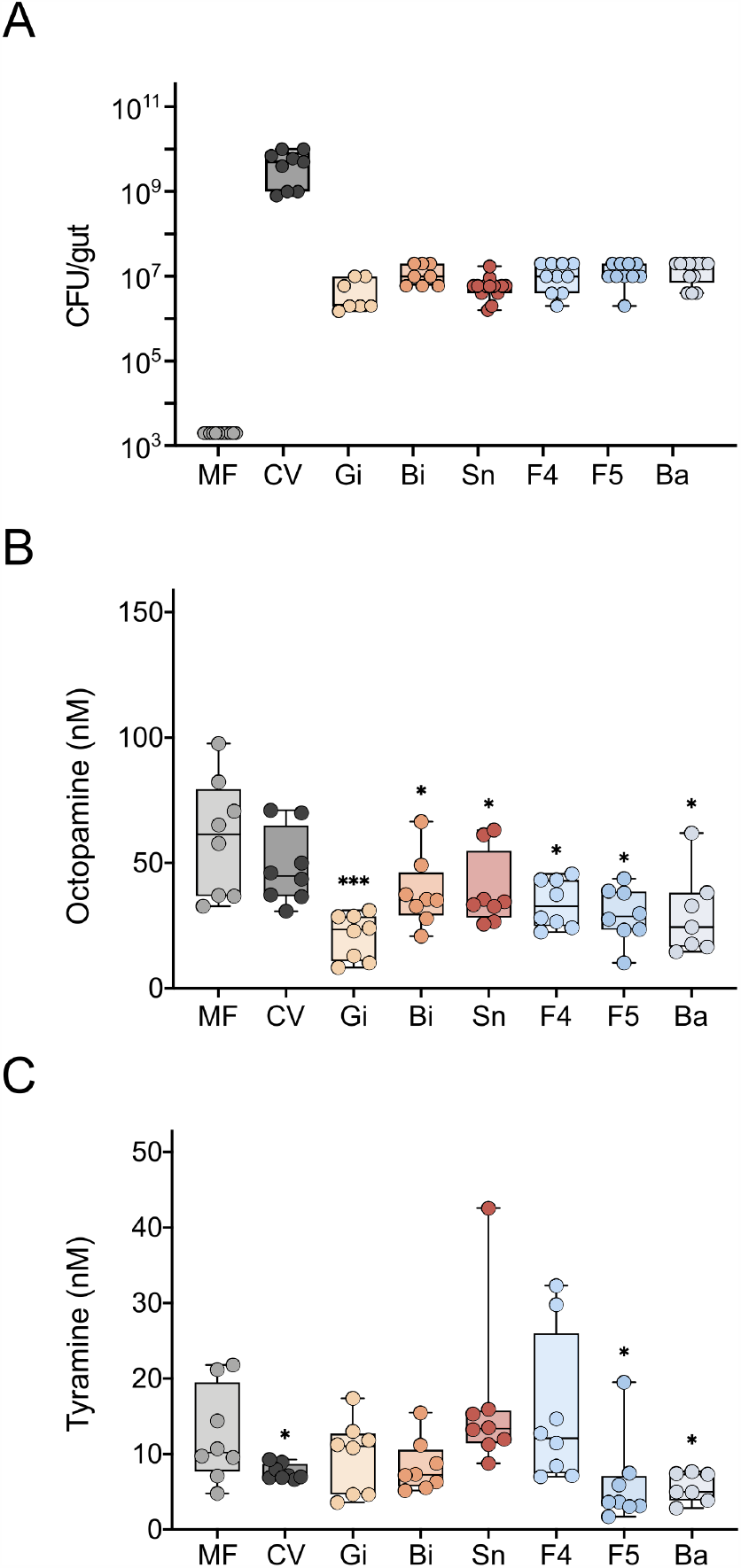
Gut microbiota impacts the concentrations of tyramine and octopamine in the honeybee brain. (**a**) Boxplots of the total CFU per gut estimated by bacteria culture for MF and mono-colonized bees, or by qPCR for the CV group. (**b-c**) Concentrations of (**b**) tyramine and (**c**) octopamine in MF (n = 8), CV (n = 8), and mono-colonized (n = 8, except n = 7 for Ba group) bee brains. Differences between bacteria-colonized bees and the MF group were tested by Mann-Whitney *u* test (*p < 0.1, **p < 0.01, ***p < 0.001). Error bars represent min and max.

**Supplementary Fig. 2.**
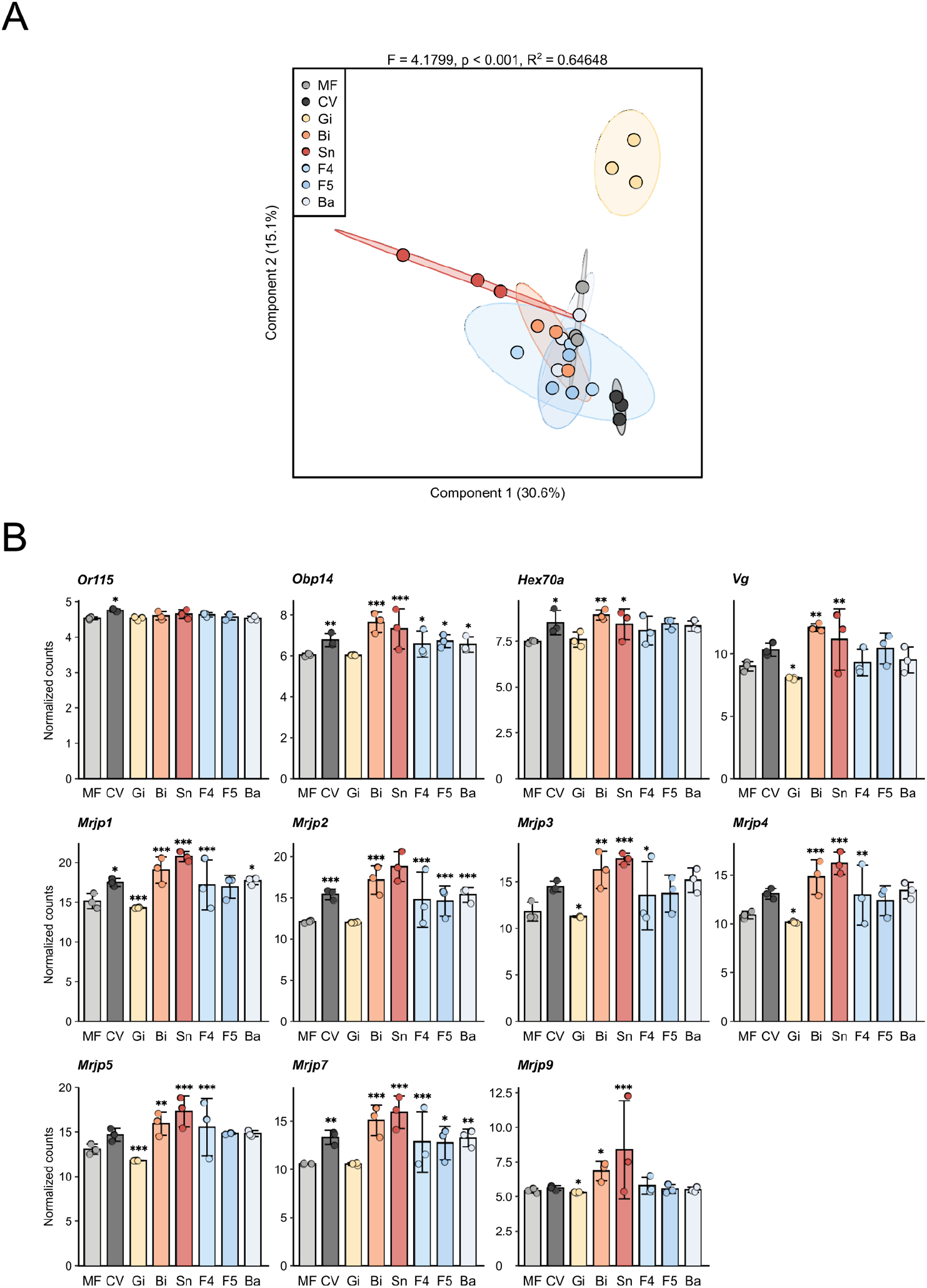
Gut microbiome impacts gene expression in the honeybee brain. (**a**) Sparse PLS-DA based on normalized gene expression in the brain of microbiota-free and bacteria-colonized bees. Group differences were tested by PERMANOVA. (**b**) Relative expression levels of differentially expressed genes in the brains of different bee groups. Differences between bacteria-colonized bees and the MF group were tested by Wald test with Benjamini-Hochberg correction (*FDR < 0.05, **FDR < 0.01, ***FDR < 0.001). Data are shown as mean ± SEM.

**Supplementary Fig. 3.**
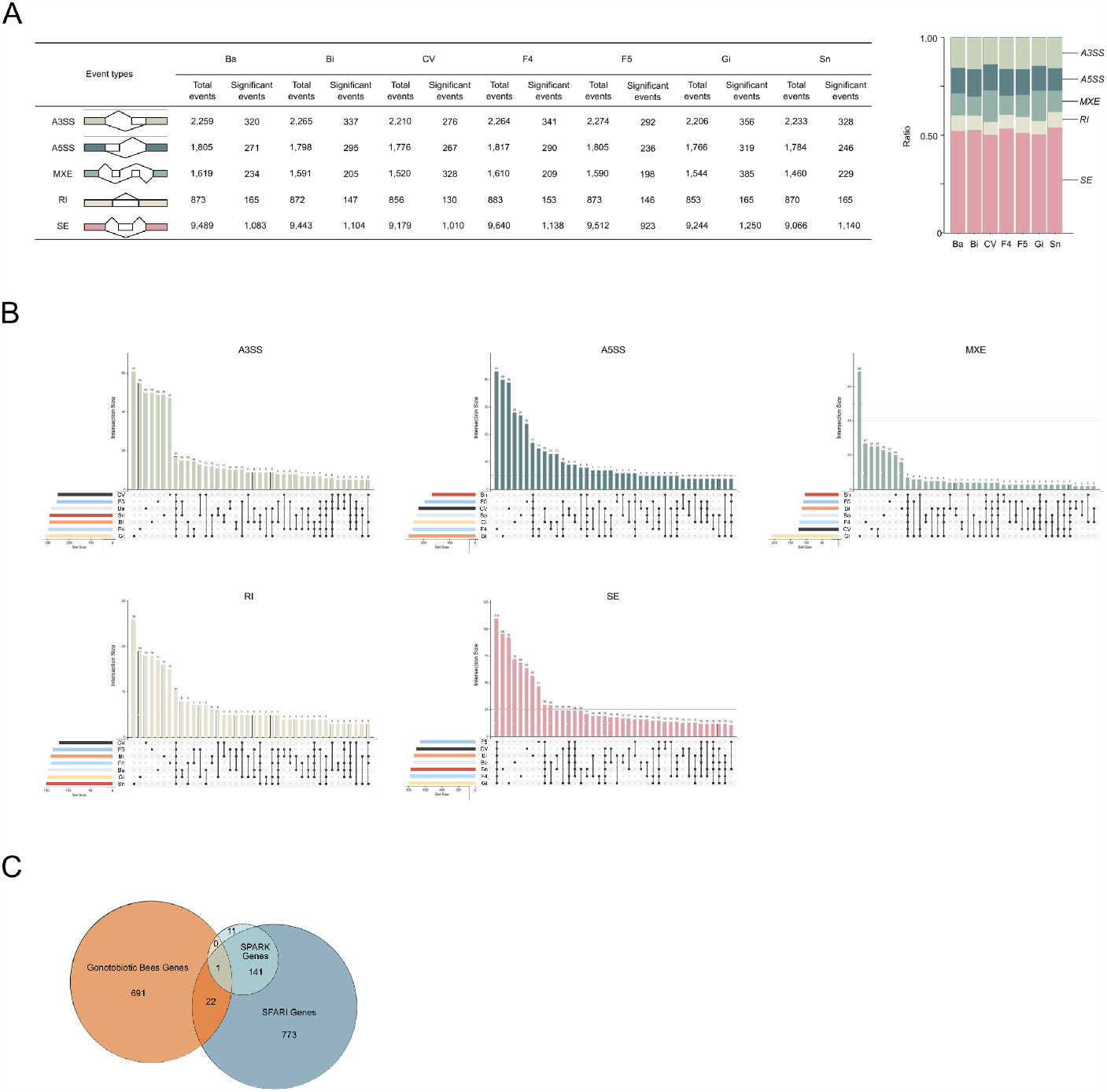
Gut microbiome impacts spliced genes in the honeybee brain. (**a**) Numbers of the differential alternative splicing events in the brains of bacteria-colonized bees compared to MF bees. Stacked column graph shows the relative abundance of different types of alternative splicing events in each group. A3SS, alternative 3’ splice site; A5SS, alternative 5’ splice site; MXE, mutually exclusive exon; RI, retained introns; SE, skipped exon. (**b**) UpSet plots showing the intersections of alternative splicing (AS) events associated with different bacteria-colonized groups. The dots and lines on the bottom right represent which intersection is shown by the bar plots above. The size of intersections is given above the bar plot. The total amount of different types of events for each bee group is given to the left of the intersection diagram. (**c**) Venn diagram of differentially expressed genes in the brains between MF and CV/mono-colonized bees (FDR < 0.05), and their overlap with the SPARK and SFARI Gene datasets. Differentially expressed genes were identified by Wald test with Benjamini-Hochberg correction.

**Supplementary Fig. 4.**
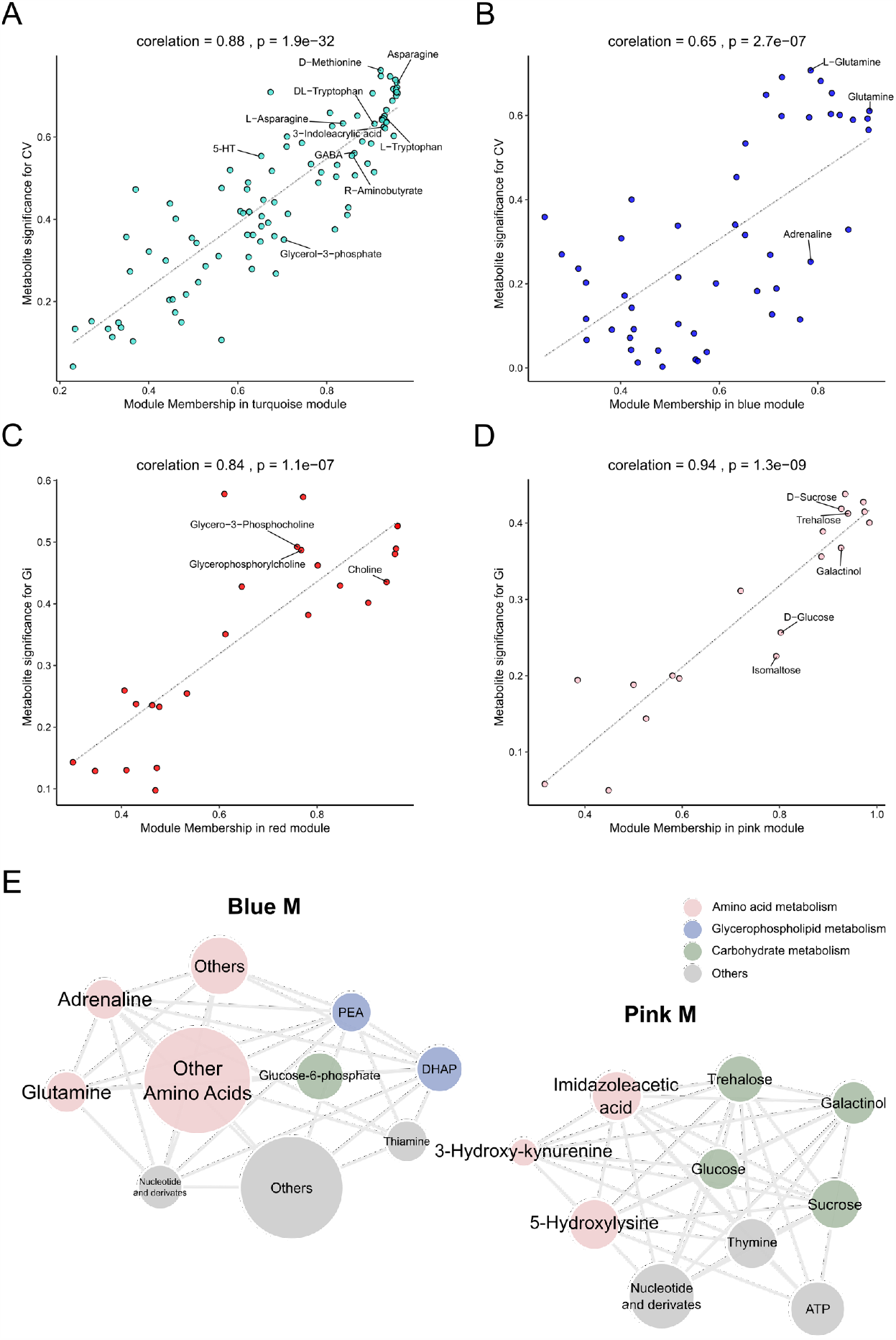
Identification of intramodular connectivity and network analysis. (**a-d**) The correlation analysis between metabolite-module connectivity (X axis) and metabolites significantly correlated with different bee groups (Y axis): (**a**) turquoise M and (**b**) blue M with CV group; (**c**) red M and (**d**) pink M with Gi group. (**e**) Network diagrams of differential metabolites in the blue and pink M. Circle colors indicate different classes of metabolites in each module, and the circle size is proportional to the total abundance of metabolites in each module.

**Supplementary Fig. 5.**
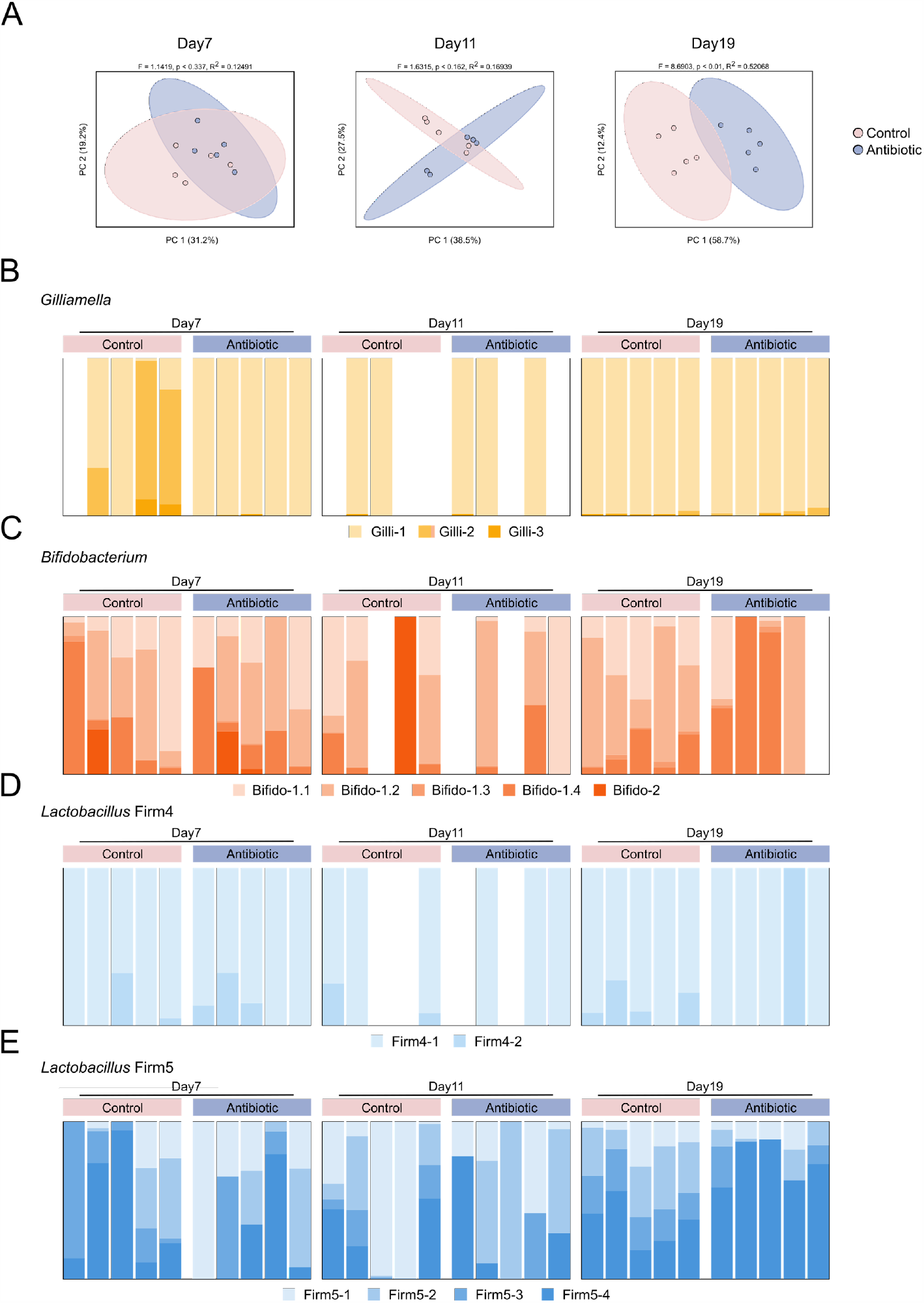
Antibiotic treatment affects the gut microbiota of honeybees. (**a**) Principal coordinate analysis of Bray-Curtis dissimilarity of gut community compositions of control and antibiotic-treated bees. Group differences were tested by PERMANOVA. (**b-e**) Compositions at SDP-level for four core bee gut members: (**b**) *Gilliamella*, (**c**) *Bifidobacterium*, (**d**) *Lactobacillus* Firm-4, and (**e**) Lactobacillus Firm-5.

**Supplementary Data 1**. Normalized gene expression levels in brains of microbiota-free and bacteria-colonized bees.

**Supplementary Data 2**. Alternative splicing events in brains of microbiota-free and bacteria-colonized bees.

**Supplementary Data 3**. Identification and biological function analysis of proteins expressed in brains of microbiota-free and conventional bees.

**Supplementary Data 4**. Raw data of all metabolites abundance in the hemolymph of microbiota-free and bacteria-colonized bees, and in the colon of antibiotic-treated and control bees.

**Supplementary Data 5**. Hemolymph metabolomic WGCNA module analysis of microbiota-free and bacteria-colonized bees.

**Supplementary Data 6**. The list of genomes of bacterial isolates in the database for MIDAS profiling.

**Supplementary Movie 1**. Olfactory learning and memory test.

## Acknowledgments

This work was founded by National Key R&D Program of China, (Grant No. 2019YFA0906500), National Natural Science Foundation of China Project 31870472.

## Author Contributions

H.Z. supervised the study; H.Z. and Z.Z. designed the study; Z.Z., Q.C. and Y.S. collected samples and performed the behavioral experiments; Z.Z. generated data and performed the data analyses with contributions from X.M. and X.H.; H.Z., Z.Z., X.M., and X.H. prepared the manuscript.

## Competing Interests

The authors declare no competing interests.

## Notes

### Competing Interest Statement

The authors have declared no competing interest.

